# A Carboxylate Switch Point Controls Long-Range Energy Transduction in Respiratory Complex I

**DOI:** 10.64898/2026.01.27.702182

**Authors:** Adel Beghiah, Patricia Saura, Terezia Kovalova, Franziska Hoeser, Thorsten Friedrich, Ville R. I. Kaila

## Abstract

Complex I is a highly intricate membrane-bound protein that powers the cellular energy metabolism by a long-range (>300 Å) proton-coupled electron transfer (PCET) reaction. Here, we investigate the unknown coupling mechanism of Complex I by probing the charge transfer reaction along its functionally central carboxylate pathway (E-channel). By combining biophysical and site-directed mutagenesis experiments with high-resolution (2.6-2.7 Å) cryo-electron microscopy (cryo-EM) and multiscale simulations, we identify a conserved carboxylate switch point (D79^NuoA^) that mediates proton transfer by establishing a kinetic gate that couples the redox chemistry to proton pumping. We find that mutation of the identified site, as found in patients suffering from severe neurodegenerative disorders, perturbs the charge transfer mechanism, and results in a drastic (>80%) reduction of the long-range PCET activity. Our combined findings illustrate mechanistic principles of molecular gates underlying long-range charge transfer reactions, and show how disease mutations perturb the function of conserved switch points in energy transduction.

---

Complex I (NADH:ubiquinone oxidoreductase) is a redox-driven proton pump that powers oxidative phosphorylation in aerobic bacteria and mitochondria^1–5^. Complex I transfers electrons from nicotinamide adenine dinucleotide (NADH, *E*_m_=-320 mV) to ubiquinone (Q, *E*_m_=+90 mV), and transduces the redox energy by pumping protons across the membrane. The proton pumping activity generates an electrochemical proton motive force (PMF) that drives active transport and the synthesis of adenosine triphosphate (ATP)^6,7^. In addition to its central role in the energy metabolism, mutations of Complex I have been linked to more than half of all known mitochondrial disorders, including severe neurodegenerative disorders such as Leigh’s syndrome^8,9^.

Complex I is a 13-45 subunit (0.5-1 MDa) L-shaped membrane-bound protein, comprising hydrophilic and membrane domains that are responsible for the respective electron and proton pumping activities (Fig. 1). The hydrophilic domain contains 8-9 iron-sulphur (FeS) centres (Fig. 1a) that mediate the 100 Å electron transfer between NADH and Q, whilst the antiporter-like subunits NuoL, NuoM, and NuoN together with the NuoA/J/K/H module of the membrane domain establish the PMF by pumping protons across the membrane (Fig. 1a). This proton-coupled electron transfer (PCET) process is highly efficient and acts across a remarkable distance of more than 300 Å. This fully reversible PCET process enables Complex I to also catalyse the reverse ΔpH-driven quinol oxidation, which results in the formation of reactive oxygen species (ROS)^10,11^, with severe physiological consequences. Despite significant structural, biochemical, and computational advances, the mechanistic principles underlying this remarkable long-range PCET process remain elusive and much debated, with several conflicting proposals in recent years^1,12–20^.

**Fig. 1:**
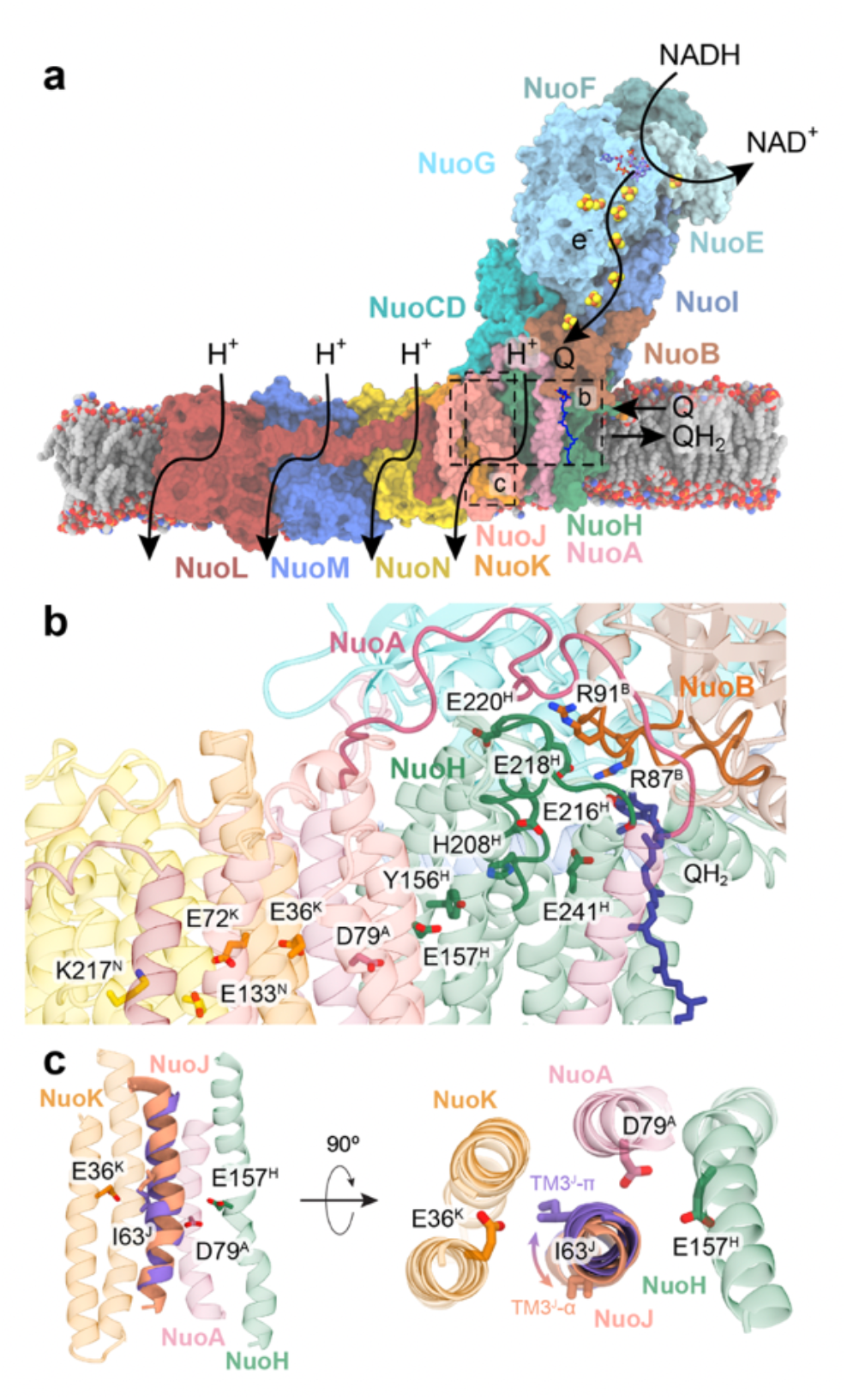
Structure and function of Complex I. **a**, *E. coli* Complex I (PDB ID: 7Z7S^13^) embedded in a model membrane. Electron transfer from NADH via the FeS clusters in the hydrophilic domain, triggers quinone reduction to quinol, which activates the translocation of four protons across the membrane domain, up to 200 Å away from the Q site. **b**, Closeup of the hinge region between the hydrophilic and membrane domains, highlighting key residues involved in the redox-driven proton pumping process, and conserved loops (here the TM5-6 loop of NuoH; PSST loop of NuoB, TM1-2 of NuoA). **c**, Redox-coupled conformational changes of TM3^J^ (π → α) favour proton transfer along the E-channel.

The proton pumping process in Complex I is initiated by the reduction of quinone to quinol (QH_2_) at the interface between the hydrophilic and membrane domains (Q site 1) that triggers both local and long-range conformational changes in the membrane domain and surrounding conserved loop regions. It was suggested^15,19,20^ that quinol diffusion into a transient Q binding site (site 2)^21^ (*cf*. also Refs.^22,23^) at the interface of NuoCD, NuoB, and NuoH, initiates a protonation cascade along the so-called E-channel region, comprising several conserved carboxylates^24^ (Fig. 1b,c). This process was also linked to the conformational switching of transmembrane (TM) helix 3 of NuoJ (TM3^J^)^12,13,20,25^ from a π-bulge form in the resting (and deactive) state of Complex I into an α-helical form, present in the active state^12,13,19,20,24–26^. According to the electrical wave propagation mechanism^1^, the α-helical TM3^J^ enables proton transfer along the E-channel towards the interface of NuoN. This, in turn, triggers charge transfer reactions in NuoN, NuoM, and NuoL, while the relaxation and back-reflection of the charge wave results in proton pumping across the membrane and finally to quinol release (see *Discussion*)^1^. In contrast, the “NuoL-only” pumping model^12,13^ assumes that the substrate protons required for the quinol formation are transferred in the opposite direction *via* the E-channel to a Q^2-^ species bound at the Q site 1, while only the terminal NuoL subunit, pumps all protons across the membrane (*cf*. also Refs.^16,27,28^ for alternative models). The “NuoL-only” model is challenged by data showing that the individual antiporter subunits^17^, as well as Complex I with deleted terminal subunits^29^, transport protons across the membrane.

Although the molecular mechanism underlying the long-range PCET process in Complex I remains much debated, all models assign a central role to the E-channel region. Key steps of this charge transfer cascade comprise water-mediated proton transfer reactions along conserved carboxylates in the E-channel that are central for the redox activity. Indeed, early biochemical studies, found that mutations of residues in the E-channel impede the NADH oxidoreductase activity of cytoplasmic membranes^30–33^, with several of these also implicated in fatal diseases such as Leigh’s syndrome^34,35^. Although these studies indirectly support that the E-channel region is functionally important, its mechanistic role remains elusive and highly debated.

To derive a molecular understanding of the redox-triggered proton transfer reactions in Complex I, we combine here biochemical and site-directed mutagenesis experiments with spectroscopic investigations, single particle cryo-electron microscopy (cryo-EM) and multiscale quantum and classical simulations, with a focus on the highly conserved D79^A^ site, which has also been implicated in the development of Leigh’s syndrome^34^. Our integrative theory-guided experimental approach provides a powerful methodology to probe the functional consequences of the highly intricate redox-driven proton pumping process in Complex I.

## Results

### Identification of a switch point that controls the long-range proton transfer in Complex I

To obtain a molecular understanding of the proton transfer reaction along the E-channel, we performed atomistic molecular dynamics (MD) simulations of the *E. coli* Complex I. To this end, we embedded the active form of the protein (PDB ID: 7Z7S^13^) into a bacterial membrane, comprising phosphatidyl ethanolamine (PE), phosphatidylglycerol (PG), and cardiolipin (CDL) (in 7:2:1 ratio), and solvated the model in a water-ion simulation box, resulting in a system with around 0.85 million atoms (Extended Data Fig. 1). During the cumulative *ca*. 4 μs MD simulations, a water wire forms between the Q site 2 along the E-channel to E157^H^, which establishes a hydrogen-bonded connection via 2-3 water molecules to D79^A^. In turn, D79^A^ connects via 4-5 water molecules to E36^K^ (Fig. 2), enabled by the α-helical conformation of TM3^J^, consistent with previous findings in *Yarrowia lipolytica* and mouse Complex I^19,20^, while the π-form of TM3^J^ disrupts this wire by the bulky I63^J^ (Extended Data Fig. 2).

**Fig. 2:**
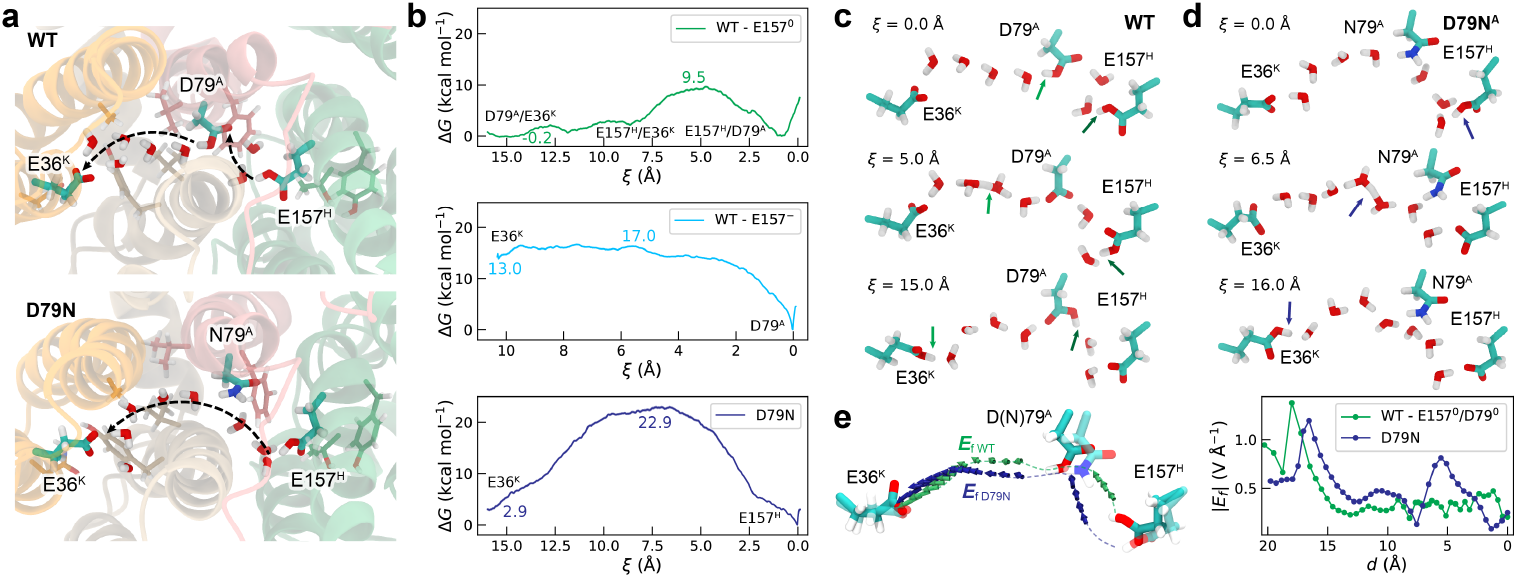
Molecular simulations of the proton transfer along the E-channel. **a**, Snapshot of the WT (*top*) and the D79N^A^ variant (*bottom*) based on QM/MM simulations. The dashed arrows indicate the proton transfer reactions. **b**, QM/MM free energy profiles of the proton transfer reactions. *Top*: WT with E157^H^ and D79^A^ protonated (green) and E157^H^ deprotonated (cyan). The proton transfer takes place from D79^A^ to E36^K^, which couples to the re-protonation of D79^A^ by E157^H^ in the former; and by direct proton transfer between D79^A^ and E36^K^ in the latter (without re-protonation of D79^A^). *Bottom*: Proton transfer in the D79N^A^ variant from E157^H^ to E36^K^ (blue). **c-d**, Intermediate structures along the proton transfer reactions in **c**, the WT and **d**, the D79N^A^ variant. The position of the transferred proton (centre of excess charge, ξ) is indicated by arrows. **e**, Electric field vectors along the proton transfer array in the WT (green) and the D79N^A^ variant (blue).

The disease-causing mutation, D79N^A^, introduces a polar headgroup along the water wire that stabilises the hydrogen-bonded network, but removes the proton-accepting D79^A^ along the array (Fig. 2a). In the D79N^A^ variant, we observe a longer proton wire, comprising 7-8 water molecules that directly connect E157^H^ with E36^K^. Interestingly, the hydration along the putative pathway is overall lower in the WT relative to the D79N^A^ variant, with the stability of the water chain correlating with the conformation of D79^A^ (Fig. 2c,d; Extended Data Fig. 3). In contrast to the WT with dynamic wetting/dewetting transitions, the D79N^A^ variant stabilises a continuous water wire along the region during the simulations (Extended Data Fig. 3). The hydration correlates with the formation of an orientated electric field along the pathway (Fig. 2e, Extended Data Fig. 4), suggesting that the proton transfer reaction could be controlled by electric field effects (cf. also refs^36,37^), which in turn are determined by the protonation states of E157^H^ and E36^K^ (Extended Data Fig. 4).

**Fig. 3:**
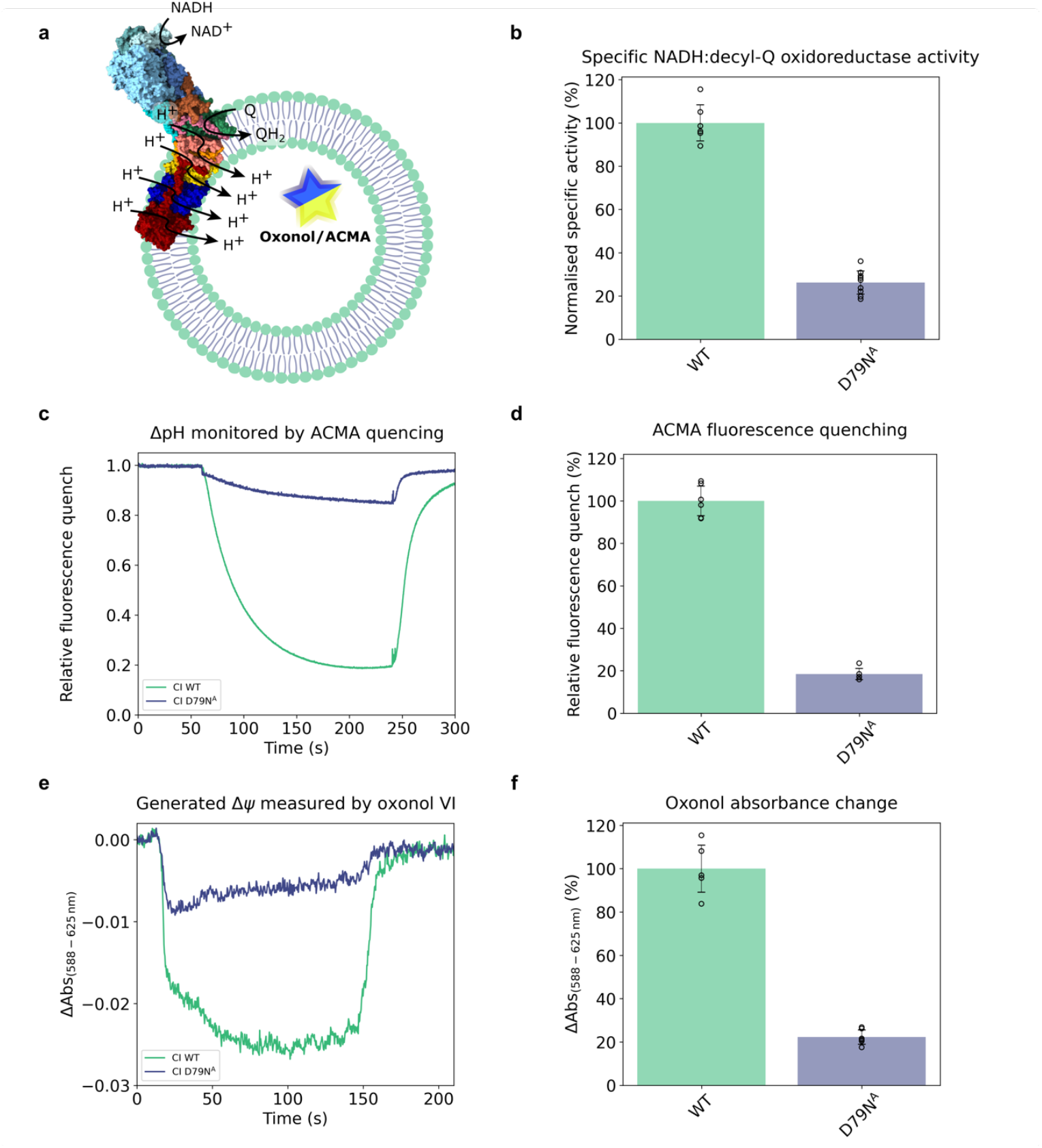
Biochemical characterisation of the WT Complex I and the D79N^A^ variant. **a**, Proteoliposome setup used for characterisation of proton pumping in Complex I using oxonol VI or ACMA. **b**, Specific NADH:DQ oxidoreductase activity (100% WT corresponding to 28.8 ± 2.41 µmol min^-1^ mg^-1^). The proton pumping activity of WT Complex I and the D79N^A^ variant assessed by following **c**, the ΔpH monitored with ACMA, and **e**, the membrane potential monitored with oxonol VI. **d, f**, Relative activity of the D79N^A^ variant based on **d**, ACMA and **f**, oxonol VI measurements. All measurements are performed in biological duplicate each tested by three technical replicates.

**Fig. 4:**
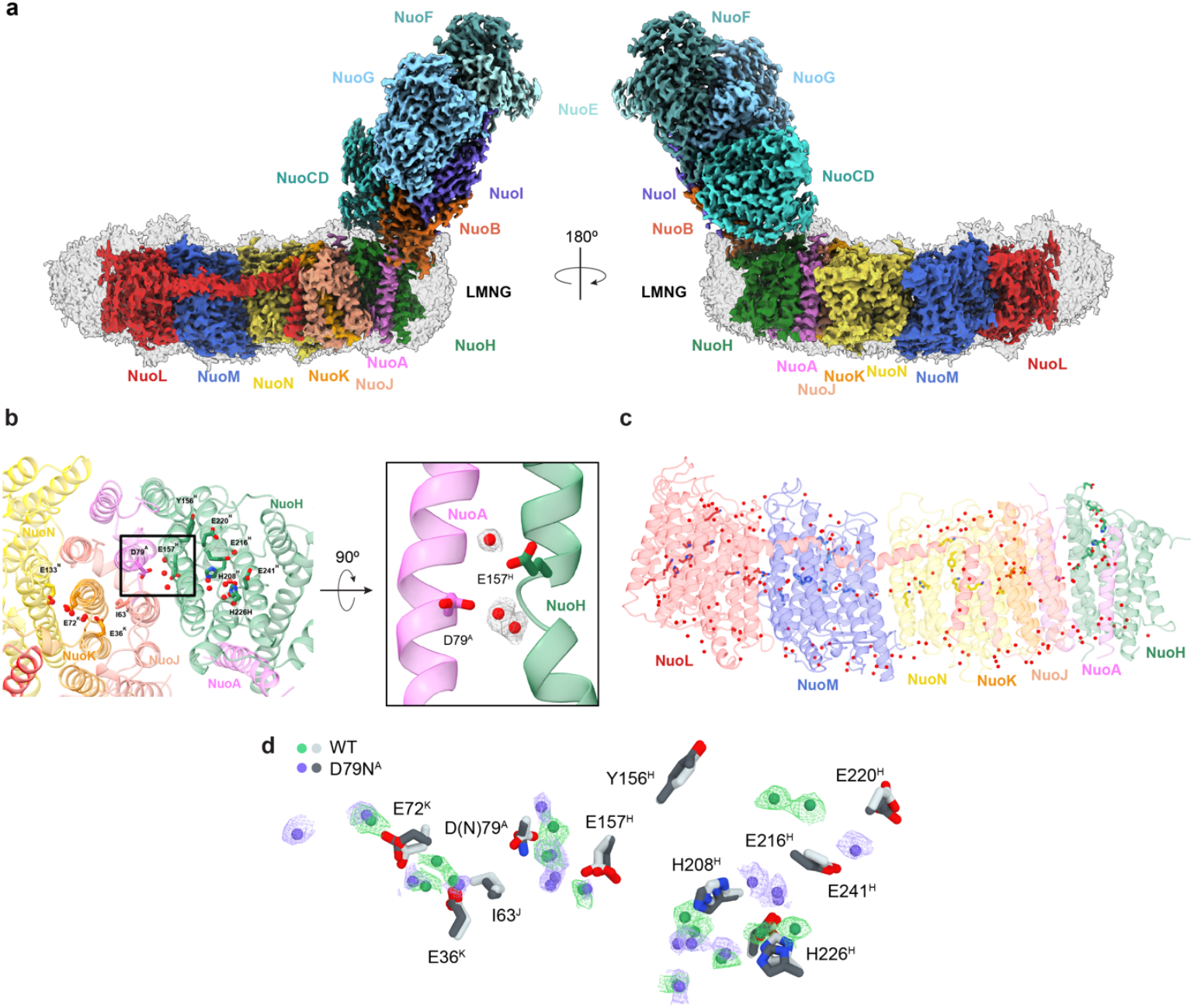
High-resolution cryo-EM structure of Complex I and the D79N^A^ variant. **a**, Cryo-EM map of the WT Complex I. **b**, Water molecules resolved in the E-channel (*inset*, side view along the membrane domain. **c**, Water molecules resolved in the membrane domain subunits **d**, Closeup of the E-channel in the WT (*green*) and D79N^A^ variant (*purple*).

To probe the proton transfer reaction along the hydrogen-bonded network, we next treated the water chain together with proton donor/acceptors (E157^H^, D79^A^, E36^K^), and their surrounding residues at the hybrid quantum/classical (QM/MM) level with hybrid density functionals, and sampled the proton transfer reaction along the wire by extensive QM/MM free energy simulations (see *Methods*). To this end, we modelled a large QM region (with *ca*. 200 atoms) to accurately describe polarisation effects at the quantum mechanical level in combination with a non-local reaction coordinate, capturing the charge propagation rather than movement of individual protons (Extended Data Fig. 1b). For the WT system, we observe a Grotthuss-type proton transfer between D79^A^ and E36^K^ that couples with a synchronous re-protonation of D79^A^ by E157^H^ (Fig. 2b, Extended Data Movie 1). The reaction has a free energy barrier of *ca*. 9.5 kcal mol^-1^ and a driving force of around -0.2 kcal mol^-1^, suggesting that the reaction takes place on μs^-1^ timescales (Fig. 2b, Extended Data Fig. 5) based on transition state theory. Interestingly, when E157^H^ is deprotonated, the free energy barrier and driving force for the D79^A^ → E36^K^ proton transfer increase to *ca*. 17 and 13 kcal mol^-1^, respectively, suggesting that the protonation state of the E-channel itself regulates the proton transfer reaction, with the origin of the barrier tuning effect arising from electrostatic effects (Fig. 2e, Extended Data Fig. 4). Our QM/MM simulations thus indicate that D79^A^ creates a kinetic gate that prevents the back-transfer of protons from E36^K^ to the former (see *Discussion*).

**Fig. 5:**
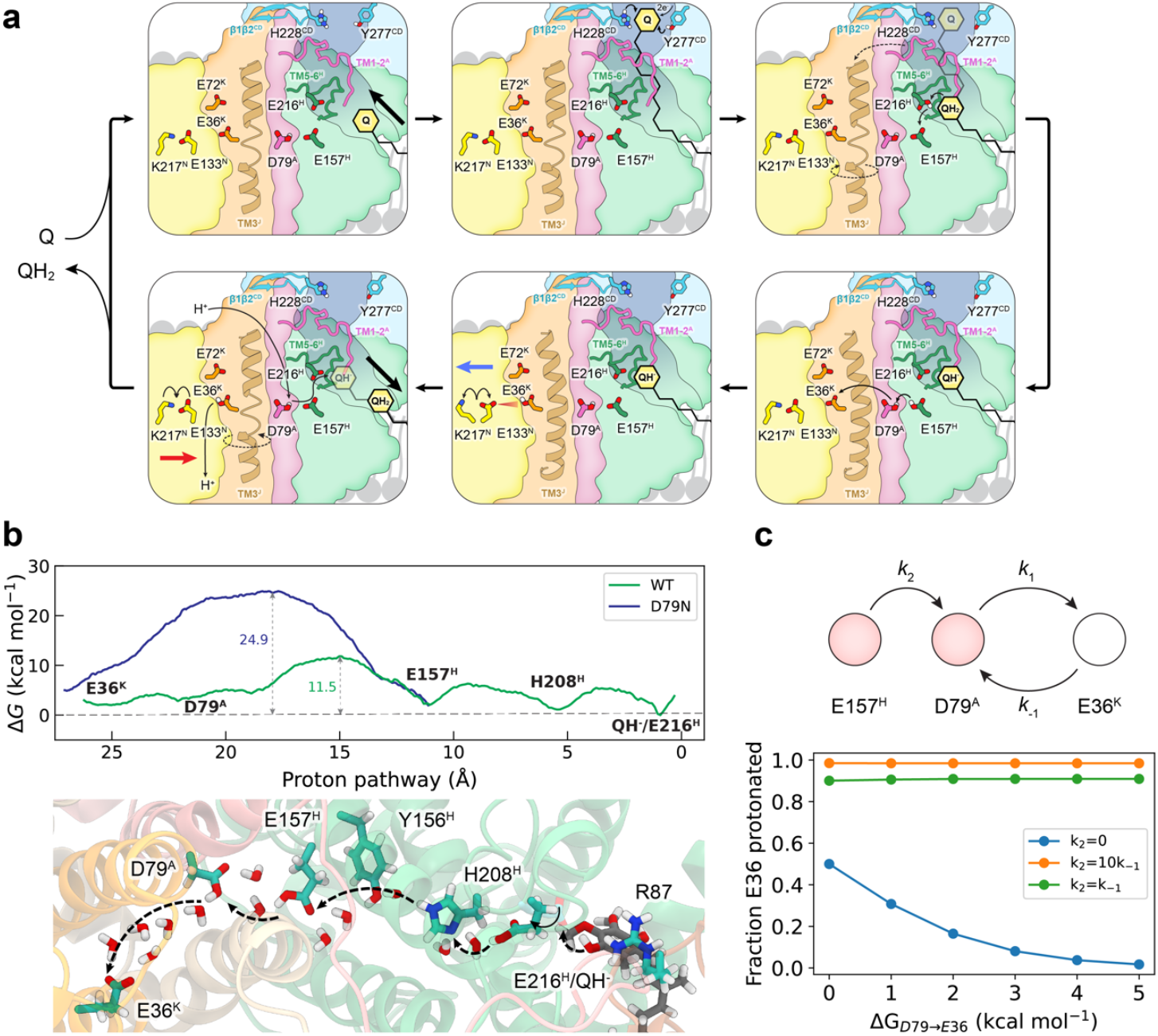
Mechanistic model of redox-coupled proton pumping in Complex I. **a**, Schematic figure of the proposed mechanism of the D79^A^ gate in mediating redox-triggered proton pumping. **b**, Energetics of the proton transfer along the E-channel. Data for the early proton transfer steps [0,10 Å] were obtained from Ref^75^. **c**, Kinetic simulations of the gating function of D79^A^ showing the fraction of E36^K^ protonation as a function of driving force and rate of the proton transfer reaction.

For the longer water chain in the D79N^A^ variant, the water-mediated proton transfer between E157^H^ and E36^K^ occurs with a significantly higher free energy barrier of *ca*. 23 kcal mol^-1^ (Fig. 2b,d, Extended Data Movie 2), thus kinetically limiting the turnover to *ca*. 100 s^-1^. This is expected to significantly decrease the population of the protonated E36^K^ as a result of the back-transfer of protons to E157^H^, thus hampering subsequent reaction cascades (see *Discussion*). Taken together, these findings suggest that D79^A^ provides a central switch point, controlling the barrier for the proton transfer reaction across the E-channel, while the D79N^A^ mutation strongly impedes PCET and results in a drastic reduction of the proton pumping and oxidoreductase activity.

### The D79N^A^ mutation perturbs the long-range PCET activity of Complex I

To experimentally assess the effects of the D79N^A^ substitution, we introduced the mutation into the pBAD_nuo_ plasmid^38^, encoding for the entire *E. coli* Complex I (see *Methods*). Expression of the *nuo*-genes from the mutated plasmid had no impact on the protein production, assembly, and stability of the complex (Extended Data Fig. 6), as also confirmed by cryo-EM structure determination (see below). We observe that the coupled NADH:O_2_ activity of cytoplasmic membranes from the mutant strain is half of that in membranes from the parent strain, as quantified with a Clark electrode (Extended Data Fig. 7, Extended Data Table 1), and demonstrating a significant effect of the mutation on electron transfer activity (cf. also Ref.^30^). To probe the effect of the mutation on the redox-driven proton pumping, we next purified the variant in the soft detergent lauryl maltose neopentyl glycol (LMNG), followed by reconstitution of the complex into proteoliposomes. The D79N^A^ variant shows a drastically diminished NADH:decylquinone (DQ) oxidoreductase activity of 26.2 ± 5.4% relative to the WT (Fig. 3b, Extended Data Table 1), as well as a drastic decrease of the proton pumping activity to 18.5 ± 2.7% relative to the WT, as quantified with the pH-sensitive fluorescence quench of 9-amino-6-chloro-2-methoxyacridine (ACMA) (Fig. 3c, d; Extended Data Table 1) in proteoliposomes. We observed a similarly large reduction in proton pumping activity of 22.3 ± 3.3% relative to the WT with the Δψ-sensitive dye oxonol VI (Figs. 3e, f; Extended Data Table 1). The proportional decrease in the electron transfer and proton pumping activities indicates that the H^+^/e^-^ stoichiometry remains unchanged in the variant, despite the drastically slower turnover. Taken together, our biochemical experiments show that the D79N^A^ substitution strongly perturbs the proton-electron coupling in Complex I, consistent with our multiscale simulations.

### Cryo-EM structure of the D79N^A^ variant reveals a perturbed hydrogen-bonding network

To assess the structural consequences of the D79N^A^ substitution, we determined the cryo-EM structure of the D79N^A^ variant and Complex I in LMNG to a resolution of 2.6-2.7 Å (Fig. 4, Extended Data Fig. 8-10). The Complex I and D79N^A^ structures are overall highly similar, showing no major structural re-arrangements (but see below), and underlining that the mutation does not impact the structural integrity of the protein. Our structures reveal more than 420 water molecules (166 in membrane domain, 258 in hydrophilic domain), 23 lipids (assigned as POPE or cardiolipin) at the interface of the antiporter subunits and around the Q entrance site, two membrane-bound Q molecules, as well as 5 Ca^2+^ ions (Fig. 4, Extended Data Fig. 10). Our WT structure resembles previously described resting state structures of the *E. coli* Complex I^13,39^, but in contrast to these, it features well-connected water molecules along the E-channel from E220^H^ to E157^H^ and D79^H^. We also observe water molecules along the central hydrophilic axis and putative proton input channels of NuoL (K305^L^, H254^L^, K342^L^ *via* H334^L^, H338^L^, K399^L^ to D400^L^), NuoM (K312^M^, D258^M^, H248^M^, K265^M^ *via* H348^M^, H322^M^ to E407^M^), and NuoN (K295^N^, Y231^N^, K247^N^ *via* H305^N^ to K395^N^), and along the E-channel (E216^H^, H226^H^, E241^H^, H208^H^) (Fig. 4b,c). The ion-pairs in NuoN (K217^N^-E133^N^, *d*=5.9 Å) and NuoM (K234^M^-E144^M^, *d*=5.8 Å) are in an open conformation, while the double ion-pair in NuoL (K229^L^-D178^L^/K229^L^-E144^L^/D178^L^-R175^L^/R175^L^/E144^L^, *d*=3.3/4.8/5.2/2.9 Å) are in their closed (salt-bridge) form, with the R175^L^ bridging halfway between E407^M^ and E144^L^, with a few water molecules bridging the residues.

In addition to the well-structured water molecules, we resolve cryo-EM density at the membrane side of NuoN that can be assigned to two Q molecules, indicating possible transient quinone interaction sites along the membrane that co-purify upon solubilisation. The determined structures also show a strong density at five unique positions in the hydrophilic domain that we assign to Ca^2+^ ions (Extended Data Fig. 10f), which is also supported by our AlphaFold3^40^ models (Extended Data Fig. 10i; *cf*. also Ref^41^). These ions are likely to be important for the structural integrity of Complex I, but they could also tune the redox potential of the FeS centres, and thus have a potential role for the electron transfer activity. Despite the overall well-resolved structure of the membrane and hydrophilic domain, the terminal edge of the Q cavity is partially open, similar to previous resting state structures^13,42^, and could arise, *e*.*g*., from interaction with the detergent upon solubilisation. Similar effects are not observed at other sites, suggesting that the determined structure accurately captures the effects of the mutation.

The membrane domain of the D79N^A^ variant, determined to a local resolution of 2.6 Å, resembles our Complex I structure, with a *root-mean-square-difference* (RMSD) of <0.6 Å (membrane domain), and comprising also more than 180 water molecules, 18 lipids, two Q molecules, and 5 Ca^2+^ ions. The D79N^A^ structure clearly resolves the substitution of the residue at position 79, while also revealing that the substitution induces some conformational changes around E220^H^, E216^H^, E241^H^, and H226^H^ near the Q site 2 (Fig. 4d), which could arise from shifted equilibrium of protonation states caused by the substitution, and supporting electrostatic interactions along the region. The conserved loops (TM1-2^A^, TM5-6^H^, PSST^B^, β1-β2^CD^) are in similar conformation in the D79N^A^ variant as in our WT structure (Extended Data Fig. 11). Moreover, although the pathway between the Q site 2 and D79^A^ contains several (*N*=17) water molecules, the region close to the D79N^A^ substitution is rather dry and features one water molecule connecting D(N)79^A^ with E36^K^ (Fig. 4) with a well-resolved π-bulge at TM3^NuoJ^ (Extended Data Fig. 10a). The TM1-2 loop of NuoA is unresolved, whilst the other conserved loops, including the TM5-6^H^, the PSST loop of NuoB, and the β1-β2 loop of NuoCD resemble previously determined resting-state structures (Extended Data Fig. 11). Taken together, our cryo-EM structures resolve high-resolution structures of Complex I and the D79N^A^ variant, revealing functional water molecules and coupling elements that provide the basis for the proton transport activity. The structural data show that the D79N^A^ substitution does not cause large structural changes, despite some specific sidechain flips along the E-channel. In particular, the D79N^A^ structure features a hydrogen-bonded connection between the Q site 2 and D79^A^, suggesting that the drastic activity changes do not result from loss of water molecules or the hydrogen-bonded network in the E-channel, but rather from perturbation in the intrinsic proton transport mechanism, as revealed by our extensive QM/MM simulations.

## Discussion

We have identified here a carboxylate switch point in the E-channel of Complex I that initiates the long-range proton pumping across the membrane. By combining molecular simulations, biochemical assays, and structural experiments, we showed that D79^N^ functions as a kinetic gate that controls the barrier for the proton transfer reaction between the quinone tunnel and the antiporter modules, while mutations of the switch point resulted in a drastic increase of the reaction barriers and drastic inhibition of the PCET reactions.

To understand the function of the identified functional element, we need to consider how the redox changes in the hydrophilic domain are linked to proton pumping across the membrane domain. Our mechanistic model proposes that the quinol formation in site 1, and the subsequent quinol binding to site 2 results in the transduction of redox energy that triggers a protonation cascade along the E-channel^19,20,43,44^. This process results in conformational changes in conserved loops around the E-channel and the Q-tunnel^13,19,20,25,45^, particularly the β1-β2 loop of NuoCD, the TM1-2 loop of NuoA, the TM5-6 loop of NuoH, and the PSST loop of NuoB that mediate the opening of the TM3^J^ gate by a π→α transition (Fig. 5a). Kim *et al*.^19^ suggested that these conformational changes enable the proton transfer along the E-channel, triggered by formation of a putative QH^-^ species at site 2^1,15,19,20,46^. A qualitatively similar reaction was recently reported^47^ based on computational studies of the mouse Complex I structure (but *cf*. also Ref. ^20^ for previous studies of the proton transfer reactions in mouse Complex I).

By combining extensive QM/MM sampling of a highly non-local reaction coordinate (centre of excess charge), we have accounted here for synchronous proton transfer reactions along the E-channel, and identified a free energy barrier for this proton transfer around D79^A^ (Δ*G*^‡^∼10 kcal mol^-1^, Fig. 5c). Our findings indicate that the proton transfer along the E-channel in the open TM3^J^ state is nearly isoenergetic, which could result in a backflow of the proton, unless kinetic gates are involved. In this regard, we suggest that the proton back-transfer is blocked by the rapid re-protonation of D79^A^ by E157^H^, with the synchronous re-protonation step increasing the back-transfer barrier by >5 kcal mol^-1^, and in turn, favouring protonation of the E36^K^/E72^K^ site. In the electrical wave propagation mechanism,^1,17,48^ the latter is a pre-requisite for the transfer of a “protonation wave” along NuoN, NuoM, and NuoL (see below). Indeed, our kinetic simulations support that the rapid re-protonation of D79^A^ fulfils such kinetic gating function by increasing the steady-state protonation of E36^K^ (Fig. 5c).

In addition to the proposed function during the forward proton transfer, D79^A^ could also serve an important role during the “back-propagation” step, during which the charge wave transfers in the opposite direction (NuoL → NuoM → NuoN → NuoA/J/K/H), and results in the ejections of pumped protons across the membrane and release of the quinol species from site 2 (Fig. 5a). We expect that a strict kinetic control of the proton transfer is central to avoid pre-mature back-transfer, which could lead to energy dissipation and RET activity. Our proton pumping experiments show that the D79N^A^ substitution drastically impedes the redox-coupled proton pumping activity, consistent with the removal of the kinetic gate at D79^A^.

The kinetic gate could thus favour the protonation of the E36^K^/E72^K^ site, which in turn induces conformational change of the K217^N^-E133^N^ ion-pair in NuoN, and triggers a lateral proton transfer towards NuoM. This lateral proton transfer results in opening of the K234^M^-E144^M^ ion-pair of NuoM, lateral proton transfer along NuoM, followed by similar conformational changes in the double ion-pair (K229^L^-D178^L^/E144^L^-R175^L^) of NuoL, mediating proton transfer and subsequent release across NuoL. The stepwise proton uptake by NuoL, relaxation of the strained ion-pair conformation, and release of the proton from NuoM follow similar steps, propagating in a reverse reaction along NuoM and NuoN to the NuoK/NuoH interface. During the back-wave, D79^A^ could control the proton transfer along the E-channel to the anionic QH^-^ species, which is released as a neutral quinol to the membrane (Fig. 5a). Our cryo-EM structure suggests that the D79N^A^ mutation also results in conformational changes near the Q site 2, supporting an allosteric crosstalk between the E-channel and the Q site 2. Together, these processes are expected to result in the relaxation of the TM3^J^ helix into its π-bulge form and re-protonation of the Y277^CD^/H228^CD^, initiating the next turnover cycle (Fig. 5a).

In contrast to the wave propagation model, where the quinol formation triggers a “forward” proton flow along the E-channel, Sazanov and co-workers^12,13^ proposed that the protons are transferred in the opposite direction along the TM3^J^ region, from E36^K^/E76^K^ via the E-channel to Q site 1. In this model, the Q reduction to Q^2-^ provides the driving force to create a charge deficit “proton hole” that results in proton uptake from the E-channel. We find that the D79N^A^ substitution does not block the formation of a water wire within the E-channel, and would thus not be expected to impede proton transfer to a doubly anionic (Q^2-^) species, if the E-channel served the function of a proton donor for the Q reduction. Accordingly, this model suggests that the proton deficient E-channel results in proton transport from NuoL *via* NuoM and NuoN, while proton uptake in NuoL and NuoM releases four protons across NuoL, although neither the directionality nor the stoichiometry cannot be linked to currently resolved structures.^12,13^ Moreover, in a variation of the NuoL-based model, Parey, *et al*.^27^ suggested that substrate protons are transferred to quinone along NuoA and NuoH upon each electron transfer step. This, in turn, was suggested to result in the uptake of two protons along the E-channel, two protons by the three antiporter subunits, and ejection of four protons from NuoL – a stoichiometry that is also difficult to rationalise based on the current structural understanding.

Our mechanism proposes that the E-channel supports a “forward-directed” proton transfer along the site, while our previous work^17^ (*cf*. also Ref.^29^) also showed that all antiporter subunits transport protons across the membrane – thus establishing strict mechanistic boundaries for the pumping model. Based on our current findings, we suggest that D79^A^ forms a kinetic gate that mediates the proton current towards the antiporter modules, while blocking the back-transfer reaction by the rapid re-protonation of D79^A^. The D79N^A^ mutation removes the kinetic gate and significantly increases the free energy barrier for proton transfer, resulting in a drastic loss of activity, possibly due to pre-mature charge-recombination reaction and slower charge flux through the gate. Remarkably, these effects take place despite the >50 Å distance between the mutation and the active site, supporting the intricate *action-at-a-distance* effects in the proton-electron coupling.

As discussed above, the D79N^A^ substitution is linked to the development of Leigh’s syndrome^34^, a fatal neurodegenerative disorder. In a case study^34^, the D79N^A^ (D66N^ND3^ in humans) was linked to a 49% reduction of Complex I activity, while the remaining respiratory chain showed normal activities. The D66N^ND3^ variant was linked to severe physiological consequences with multiple metabolic abnormalities, although the molecular principles resulting in these effects remained unknown. These severe physiological effects could result from charge recombination due to the perturbed proton gate in the E-channel, which in turn is expected to dissipate energy, possibly enhancing reverse electron transfer and production of ROS^10^.

In summary, our combined findings reveal that D79^A^ functions as a central switch point, mediating directional proton transfer along the E-channel. By combining multiscale quantum and classical simulations with site-directed mutagenesis, electron transfer assays, proton pumping experiments in proteoliposomes, and high-resolution cryo-EM, we find that the kinetically efficient de- and re-protonation of D79^A^ prevents a premature back-flow during the charge propagation process. The D79N^A^ mutation resulted in a significant increase of the proton transfer barrier based on our hybrid quantum/classical (QM/MM) free energy simulations, consistent with the drastic reduction of the coupled electron and proton transfer activities observed in our proteoliposome experiments. Moreover, our cryo-EM experiments showed that the substitution resulted in some conformational changes of titratable residues near the transient Q binding site (site 2), indicative of electrostatic coupling effects within the E-channel, but not in a disassembly of the water-mediated contacts within the channel. Taken together, our findings highlight the importance of molecular gates in mediating the long-range proton-coupled electron transfer in Complex I, and provide a molecular basis for understanding the mechanism underlying how mutations result in the development of severe diseases.

## Methods

### Mutagenesis and cloning

The pBAD_nuo_ plasmid, encoding for entire *E. coli* Complex I, was used as a template for site-directed mutagenesis. The D79N^A^ substitution was introduced via QuickChange using a pair of overlapping oligonucleotides (Extended Data Table 2). The PCR was carried out with the KOD Hot Start DNA polymerase (Novagen). Following amplification, the PCR product was digested with DpnI and the DH5α_Δnuo_ cloning strain (F-Φ80*lac*Z*Δ*M15 *Δ*(*lac*ZY A-*arg*F) U169 *recA1 endA1 hsdR17* (rk-mk+) *gal-phoA supE44* λ-*thi-1 gyrA96 relA1, Δnuo*)^49^ was transformed with the modified plasmid. Transformants were selected on chloramphenicol plate and single colonies were grown overnight for plasmid DNA isolation. The mutation was confirmed by Sanger sequencing (GATC Eurofins, Konstanz, Germany).

### Bacterial growth and membrane isolation

The BW25113_*Δndh nuo::nptII_FRT*_ expression strain^38,50^ was transformed with the native and modified pBAD_nuo_ plasmid. A single colony was inoculated into LB medium in the presence of 30 µg mL^-1^ chloramphenicol and grown overnight at 37°C, 180 rpm. The overnight pre-culture was used to inoculate (50 mL per L of medium) into Complex I autoinduction media^51^ (1% (w/v) peptone, 0.5% (w/v) yeast extract, 0.4% glycerol, 25 mM Na_2_HPO_4_, 25 mM KH_2_PO_4_, 50 mM NH_4_Cl, 5 mM Na_2_SO_4_·10H_2_O, 2 mM MgSO_4_·7H_2_O, 0.2% (w/v) L-arabinose, 0.05% (w/v) glucose, 30 mg per L Fe-NH_4_-citrate, 0.5 mM l-cysteine, 50 mg per L riboflavin), supplemented with 30 µg mL^-1^ chloramphenicol for selection. Cells were harvested at OD_600_ = 4.0, flash frozen, and stored at -80°C or resuspended in cell resuspension buffer in a 3:1 ratio (v:w). The resuspension was supplemented with 0.1 mg mL^-1^ DNase and 0.5 mM 4-(2-aminoethyl) benzenesulfonyl fluoride hydrochloride (AEBSF) for protease inhibition. Cells were lysed by three cycles through an Emulsiflex at 15,000 psi. Cell debris were removed by centrifugation (10,000 g, 20 min, 4°C) followed by an ultra-centrifugation (180,000 g, 1.5 h, 4°C) to collect cytoplasmic membranes.

### Protein purification

Cytoplasmic membranes were resuspended in a 1:3 (w:v) ratio in membrane resuspension buffer before solubilisation with 2% lauryl maltose neopentyl glycol (LMNG, BioNordika), for 1 h at room temperature under slow stirring. Solubilised membrane proteins were collected as supernatant after ultra-centrifugation (180,000 g, 30 min, 4°C). The sample was supplemented with 20 mM imidazole, prior to loading onto a 24 mL Ni-IDA column for immobilised metal affinity chromatography (IMAC). The column was washed first with buffer A (Extended Data Table 3), followed by a second wash step with 20% buffer B. *E. coli* Complex I was eluted using 60% buffer B (Extended Data Table 3), and relevant fractions were pooled and concentrated to 1 mL for size exclusion chromatography (SEC, HiLoad 16/600 Superose 6 prep grade, 1 mL min^-1^) using buffer C (Extended Data Table 3).

### Protein reconstitution

*E. coli* polar lipids (ECPL; Avanti, 25 mg mL^-1^ in CHCl_3_) were dried under N_2_ (g) stream, and desiccated overnight. The lipid film was resuspended at 5 mg mL^-1^ in reconstitution buffer (Extended Data Table 3). To allow protein incorporation, 0.4% of sodium cholate was added to partially solubilise the liposome bilayer, followed by addition of 100 µg of WT Complex I or the D79N^A^ variant. The mixture was incubated for 30 min at 4°C. Detergent and non-reconstituted protein were removed by pre-equilibrated PD-10 desalting column (Cytiva) in reconstitution buffer (Extended Data Table 3). The eluate was sedimented by ultra-centrifugation (180,000 g, 30 min, 4°C) and resuspended in 50 µL in the reconstitution buffer.

### Activity assays

#### NADH:O_2_ oxidoreductase activity

The NADH:O_2_ oxidoreductase activity was measured using a Clark electrode. Cytoplasmic membranes (5 µL; 48-62 mg mL^-1^) were incubated at 30°C, in a final volume of 1 mL of membrane resuspension buffer for 2 minutes. Once a stable baseline was established, the reaction was initiated by adding 0.4 mM NADH until complete oxygen consumption.

#### NADH: DecylQ oxidoreductase activity

The NADH:DecylQ oxidoreductase activity was monitored at 340 nm. 1 µL of *Ec*CI was incubated together with bo3 oxidase and 100 µM DecylQ, in 1 mL final volume of reconstitution buffer. The measurement was initiated with 100 µM NADH for two minutes, at 30°C.

#### Ferricyanide assay

The orientation of Complex I in proteoliposomes (PL) was assessed by monitoring the K_3_Fe(CN)_6_ oxidation rate at 420 nm over time. The assay was performed by incubating 0.2 mM NADH in 1 mL of FeCN at 30°C. Once a stable baseline was established, 1 µL of PL was added, followed by addition of DDM to disrupt the PL, enabling the population of *inward*-oriented Complex I to consume NADH. The fraction of *outward*-facing to total Complex I was determined, with the ratio providing the percentage of *outward* facing proteins. The PL volume was carefully adjusted to ensure an equal number of *outwards*-facing proteins during proton pumping experiments.

#### Proton pumping activity – detection of electrical gradient using oxonol-VI

The generated Δψ of Complex I or the D79N^A^ variant reconstituted into proteoliposomes was assessed using the oxonol buffer by monitoring the absorbance difference between 588 nm and 625 nm. Proteoliposomes with Complex I (5 µL) or adjusted volume for the variant, were incubated with 160 µM decylubiquinone (DQ) in DMSO, to a final volume of 1 mL oxonol buffer at 30°C (Extended Data Table 3). The sample was equilibrated for 3 minutes before initiating the reaction with 100 µM NADH after 15 s of baseline measurement. The PMF was dissipated after 2 min by an addition of 4 µg mL^- -1^ gramicidin.

#### Proton pumping activity – detection of pH gradient using ACMA

The proton pumping activity of Complex I or the D79N^A^ variant reconstituted into proteoliposomes was assessed using the ACMA (9-amino-6-chloro-2-methoxyacridine) buffer (Extended Data Table 3), by monitoring the ACMA fluorescence quench at 480 nm upon excitation at 410 nm. 5 µL of proteoliposomes with Complex I were used for characterisation of the proton pumping together with 100 µM DQ. The sample was equilibrated for 5 min at 30°C. After one minute of baseline equilibration, the reaction was initiated by an addition of 100 µM NADH, for 3 min. The ΔpH was dissipated by an addition was dissipated by addition of 4 µg mL^-1^ gramicidin.

### Cryo-EM sample preparation

Purified protein sample was loaded on a size-exclusion chromatography column (HiLoad 16/600 Superose 6 prep grade) in 50 mM MES pH 6.0, 150 (dataset 2) or 300 mM KCl (dataset 1), 5 mM MgCl_2_, 2% glycerol, and 0.005% LMNG for Complex I WT and 50 mM MES pH 6.0, 150 mM KCl, 5 mM MgCl_2_, 2% glycerol, and 0.005% LMNG for the D79N^A^ variant. The sample was concentrated to 4 mg mL^-1^ and mixed with 0.25 mg mL^-1^ *E. coli* polar lipids (Avanti) and 0.1% CHAPS final concentration prior to blotting on ANT grids (ANTcryo ANTA grids, Cu300-R1.2/1.3). Grids were first glow discharged at 20 mA for 120 s (PELCO easiGlow), before applying 3 µL sample on the grid and blotting for 2 s at 4°C (100% humidity, blot force 0), followed by plunge freezing in liquid ethane, using a Vitrobot Mark IV (Thermo Fisher Scientific).

### Cryo-EM data collection

Images were collected using a Titan Krios G3i electron microscope (300 kV, Thermo Fisher Scientific) equipped with K3 Gatan detector. The WT Complex I (13,882 + 5,509 exposures) and the D79N^A^ datasets (10,621 exposures), were recorded in electron-counting mode at a nominal magnification of 105,000 (0.825 Å pixel^-1^), with a camera exposure rate of 14/14.6/13.5 e^-^ pixel^-1^ s^-1^ and total exposure of 1.95/1.86/2.02 s, respectively (Extended Data Table 4).

### Cryo-EM data processing and structure refinement

The datasets for Complex I and the D79N^A^ variant were processed using cryoSPARC v4.4.0^52^. Micrographs were imported, followed by patch motion correction and patch CTF estimation. The WT structure was solved from micrographs from two different datasets, with dataset 1 ([KCl]=300 mM) comprising 13,882 micrographs, and dataset 2 ([KCl] =150 mM) with 5,509 micrographs. Both datasets were processed separately following the same workflow (Extended Data Fig. 8), which consisted of motion correction, CTF estimation, and template generation using 2000 micrographs, and a Blob picker job. Generated template 2D classes were used for template picker job, and resulted in the extraction of 3,961,114 particles for dataset 1 and 1,343,714 particles for dataset 2. The particle sets were reduced by multiple rounds of 2D and *ab initio* classification to a final number of particles of 63,982 (dataset 1) and 46,649 particles (dataset 2). Final particles from dataset 2 were exported and combined with dataset 1 to yield 110,621 particles. Combined particles were separated with 2D classification into 90,534 particles, and then merged to generate two classes with 70,502 particles, used for non-uniform refinement to solve the Complex I structure to an overall resolution of 3.0 Å, while the membrane domain and hydrophilic arm were resolved in local refinement to a resolution of 2.8 Å and 2.6 Å, respectively. The locally refined models were manually refined with Coot 0.9.8.3,^53^ followed by real-space refinement using Phenix 1.20.1^54^. The models were validated with MolProbity^55^ and combined to generate a composite map that was refined using Coot^53^ and Phenix^54^.

For the D79N^A^ dataset, particles were template picked, resulting in 3,342,145 selected particles. Multiple 2D and *ab initio* classifications narrowed down the number of particles to 192,869. Heterogeneous refinement was used to select 86,927 particles, followed by non-uniform refinement, resulting in particles refined at 2.9 Å (Extended Data Fig. 8-9, Extended Data Table 4). To better resolve the hydrophilic and membrane domains, local motion correction and local refinement using appropriate masks were used, resulting in final local refined maps with respective resolution of 2.5 Å and 2.6 Å (Extended Data Fig. 6). The locally refined models were manually refined with Coot 0.9.8.3,^53^ followed by real-space refinement using Phenix 1.21.2^54^. The models were validated with MolProbity^55^.

### Atomistic MD simulations

Atomistic molecular dynamics (MD) simulations were performed to probe the dynamics along the E-channel in the WT Complex I and the D79N^A^ variant. The initial coordinates of the protein were obtained from a cryo-EM structure of *E. coli* Complex I (PDB ID: 7Z7S^13^), with the TM3^J^ in an α-helical conformation. The missing parts of the protein were modelled using the loop modelling feature in Modeller. Protonation states were initially assigned with PropKa3^56^, and different protonation states along the E-channel were sampled to the protonation-state dynamics on microsecond timescales (Extended Data Table 5). The protein was embedded in a lipid membrane mimicking the composition of the *E. coli* polar lipid, with 67% phosphatidyl-ethanolamine (POPE), 23% phosphatidyl-glycerol (POPG), and 9.8% cardiolipin (CDL). Lipid molecules resolved in the cryo-EM structure were additionally incorporated in the model, and modelled as POPE. The membrane was created with CHARMM-GUI^57^. The membrane-protein system was solvated with TIP3P water molecules and 150 mM NaCl concentration, resulting in systems with around 850,000 atoms. The CHARMM36 force field^58^ was used to describe protein, solvent, and membrane molecules, whereas DFT-derived in-house parameters were employed for the cofactors. Simulations were propagated in an *NPT* ensemble, with *T* = 310 K and *p* = 1 bar, using an integration time step of 2 fs. Long-range electrostatic interactions were described by the Particle Mesh Ewald approach. All MD simulations were performed using NAMD3.06b and NAMD2.14.^59^ Analysis and visualisation were performed with VMD^60^, ChimeraX^61^, and MDAnalysis^62,63^. See Extended Data Fig. 1 and Extended Data Table 5 for further details.

### QM/MM free energy calculations

The energetics of the proton transfer reactions along the E-channel were explored by hybrid quantum/classical (QM/MM) free energy calculations. To this end, classically relaxed snapshots were selected from the MD trajectories (simulation S1 and S4 for WT, and simulation S5 for the D79N^A^ variant, Extended Data Table 6). The QM/MM system included subunits NuoH, NuoA, NuoK, NuoJ, NuoN, and part of NuoB and NuoCD, together with the surrounding membrane, water molecules and ions. The boundary between the QM region and the MM region was described by a link atom scheme, introduced between Cα and Cβ atoms of protein residues. The QM region comprised the sidechain of D79^A^/N79^A^, E36^K^, E157^H^, F75^A^, V76^A^, T153^H^, Y156^H^, V58^J^, Y59^J^, A62^J^, N40^K^, and A73^K^, together with 23-26 water molecules, resulting in a QM region of *ca*. 200 atoms (Extended Data Fig. 1b). An active region of 10 Å around the QM region was allowed to relax during QM/MM simulations, while keeping the rest of the system fixed. QM atoms were treated at the B3LYP-D3/def2-SVP level of theory^64–67^, while classical atoms were described by the CHARMM36 force field^58^.

The free energy profiles were explored by the umbrella sampling (US) method. In this regard, the initial positions of the US windows were generated by defining a linear combination of bond-breaking and bond-forming distances from the proton donor to the acceptor,

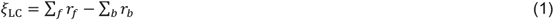

where *r*_*f*_ refers to the distance of the forming bonds, and *r*_*b*_ to the distance of the breaking bonds. This reaction coordinate was projected onto a modified centre of excess charge reaction coordinate^68^, which projects the centre of excess charge vector positions onto the donor-acceptor vector,

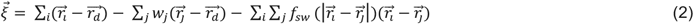

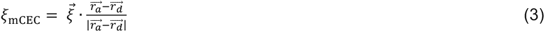

where 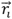refers to the position of hydrogens, 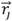to the position of heavy atoms, 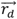 to the proton donor, and 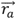 to the proton acceptor positions, *w*_*j*_corresponds to a weight associated with every heavy atom *j*, representing the minimum number of protons bound to this atom in its reference state, and *f*_*sw*_ is a switching function,

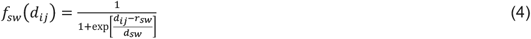

with *r*_*sw*_ = 1.3 Å and *d*_*sw*_ = 0.05 Å. Each window was sampled along the mCEC reaction coordinate using a harmonic potential with a force constant of 100 kcal mol^-1^ Å^-2^, and with a 0.25 Å separation between consecutive windows. Each window was propagated at *T*=310 K for 6-8 ps, with a 0.5 fs integration timestep, resulting in a sampling of 250-520 ps per free energy profile (around 1200 ps in total, Extended Data Table 6). The free energy profiles were derived by the weighted histogram analysis method (WHAM)^69^, as well as by the multistate Bennett acceptance ratio (MBAR) methods^70^ (Extended data Figure 5). All QM/MM sampling were performed with the FermiONs++^71^.

### Hydration analysis

Analysis of the water occupancy in the region connecting E157^H^ with E36^K^ was performed based on classical MD simulations. In this regard, a pathway was identified by a tunnel analysis, as implemented in CAVER^72^, using a probe radius of 0.9 Å and starting coordinates at the position of Y156^H^, while the sidechains of E157^H^, D79^A^, and E36^K^ were excluded from the cavity search. Water molecules along the identified pathway (highest cluster population) were analysed along the MD trajectories, using a 2 Å radius centred on the tunnel coordinates of the region between D(N)79^A^ and E36^K^. The hydration percentage was calculated based on the ratio of frames containing water molecules and the total number of frames in the MD trajectory (hydration % = *N*_occupied_/*N*_total_×100). See Extended Data Fig. 1 and Extended Data Table 6 for further details.

### Electric field analysis

Electric fields were computed along the pathway connecting E157^H^ via D(N)79^A^ to E36^K^ based on the classical MD simulations. NuoA, NuoH, NuoJ and NuoK were included in the calculations, while solvent molecules were excluded. The electric fields were calculated at each tunnel point by averaging 1 frame/ns from the two simulation replicas for each condition (Extended Data Fig. 4). For each MD frame, the electric field at each point was calculated using,

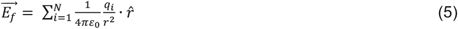

where *q*_*i*_ are the atomic partial charges of the surrounding system, and *r* is the distance to each point charge, while charges within 2 Å of the probe position were excluded from the analysis. The electric field calculations were performed with TUPA^73^, while the fields were visualised in VMD^60^ and MDAnalysis^62,63^.

### Kinetic master equations simulations

The proton transfer kinetics were modelled by a kinetic master equation model (Fig. 5c) of the proton transfer reaction based on,

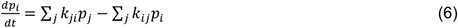

The elementary proton transfer reactions were modelled by screening different driving forces, ΔG = *k*_D79→E36_ /*k*_E36→D79_, by different ratio of rates (*k*_E157→D79_ = 0 s^-1^, *k*_E36→D79,_ 10*k*_E36→D79_, with *k*_D79→E36_ = *k*_E36→D79_ ∼(1 μs)^-1^ based on transition state theory. The master equations were integrated numerically using COPASI^74^.

## Acknowledgments

This work was supported by the Knut and Alice Wallenberg Foundation (2019.0251 and 2024.0220 to V.R.I.K), the Göran Gustafsson Foundation for Research in Natural Sciences and Medicine (to V.R.I.K.), by the Swedish Research Council (2020-04081 and 2025-04607 to V.R.I.K.), and by the Deutsche Forschungsgemeinschaft (DFG GRK2202; 235777276/RTG & SPP1927; FR 1140/11-2 to T.F.). Computing resources were provided by the National Academic Infrastructure for Supercomputing in Sweden (NAISS 2025/1-33, 2024/1-28). The cryo-EM data were collected at the Cryo-EM Swedish National Facility funded by the Knut and Alice Wallenberg, Family Erling Persson and Kempe Foundations, SciLifeLab, Stockholm University and Umeå University.

## Data availability

Data supporting the findings of this manuscript are available from the corresponding authors upon reasonable request. A reporting summary for this Article is available as a Supplementary Information file. The source data underlying Figs. 1-4 and Extended Data Figs. 1-11 are provided as a Source Data file. Model and cryo-EM density maps of the WT and D79N^A^ variant Complex I were deposited to the PDB under accession codes 9TAJ, 9TAK, 9TAL, 9TAM, 9TAN, and 9TAO and the EM Data Bank under accession codes EMD-55748, EMD-55749, EMD-55750, EMD-55751, EMD-55752, and EMD-55753, respectively.

## Author contributions

V.R.I.K. designed the study and directed the project;

A.B. and F.H. constructed, isolated, and experimentally characterised the protein;

A.B. performed biophysical proton pumping assays and activity measurements;

P.S. performed multiscale simulations;

A.B. and T.K. collected cryo-EM data;

A.B. and T.K. refined cryo-EM models;

A.B., T.K., P.S., V.R.I.K. analysed cryo-EM models;

A.B., P.S., T.K., F.H., T.F., V.R.I.K. analysed data;

A.B., P.S., V.R.I.K. wrote the manuscript with input from all authors.

## Additional information

Supplementary information is available for this paper at XXX. Correspondence and requests for materials should be addressed to V.R.I.K.

