## Supplementary Information for "A Carboxylate Switch Point Controls Long-Range Energy Transduction in Respiratory Complex I"

**Content**

**Extended Data Fig. 1.** Molecular simulation setups.
**Extended Data Fig. 2.** Disruption of water wire along the E-channel in the resting state.
**Extended Data Fig. 3.** Hydration dynamics along the E-channel in Complex I and the D79N$^{A}$ variant.
**Extended Data Fig. 4.** Electric field effects along the E-channel.
**Extended Data Fig. 5.** Convergence of the QM/MM free energy simulations.
**Extended Data Fig. 6.** Protein purification.
**Extended Data Fig. 7.** Oxygen consumption in cytoplasmic membranes.
**Extended Data Fig. 8.** Cryo-EM data analysis and validation of WT Complex I.
**Extended Data Fig. 9.** Cryo-EM data analysis and validation of the D79N$^{A}$ variant.
**Extended Data Fig. 10.** Example cryo-EM densities of key regions.
**Extended Data Fig. 11.** Structure of conserved loops.

**Extended Data Table 1 |** Activity of WT Complex I and the D79N$^{A}$ variant.
**Extended Data Table 2 |** List of designed primers.
**Extended Data Table 3 |** List of employed buffers.
**Extended Data Table 4 |** Cryo-EM data collection, refinement, and validation statistics.
**Extended Data Table 5 |** List of MD simulations.
**Extended Data Table 6 |** List of QM/MM simulations.

**Extended Data Movie 1 |** Proton transfer along the E-channel of WT Complex I.
**Extended Data Movie 2 |** Proton transfer along the E-channel of the D79N$^{A}$ variant.

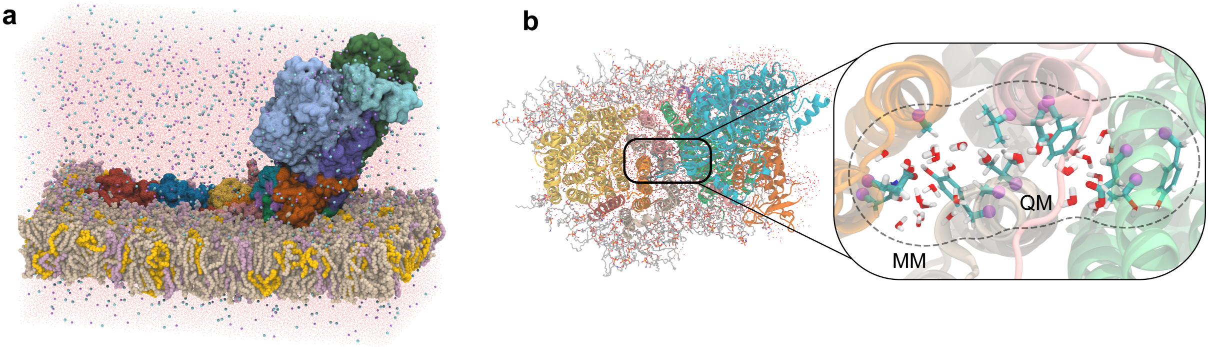


**Extended Data Fig. 1 | Molecular simulation setups. a**, Setup of the classical MD simulations. **b**, Setup of the QM/MM calculations (*left*), and a closeup of the *ca.* 200 atoms in the QM region (*inset*, *right*). Link atoms are shown as purple spheres.

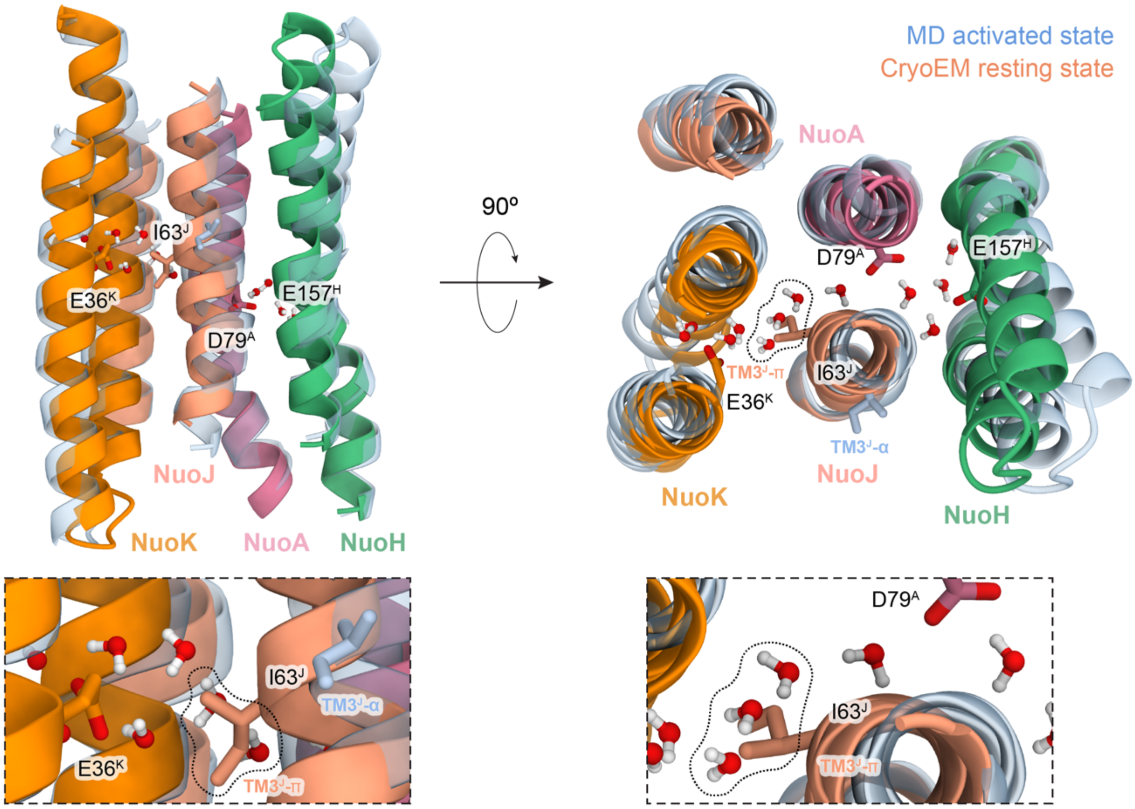


**Extended Data Fig. 2.** Disruption of the water wire along the E-channel in the resting state.

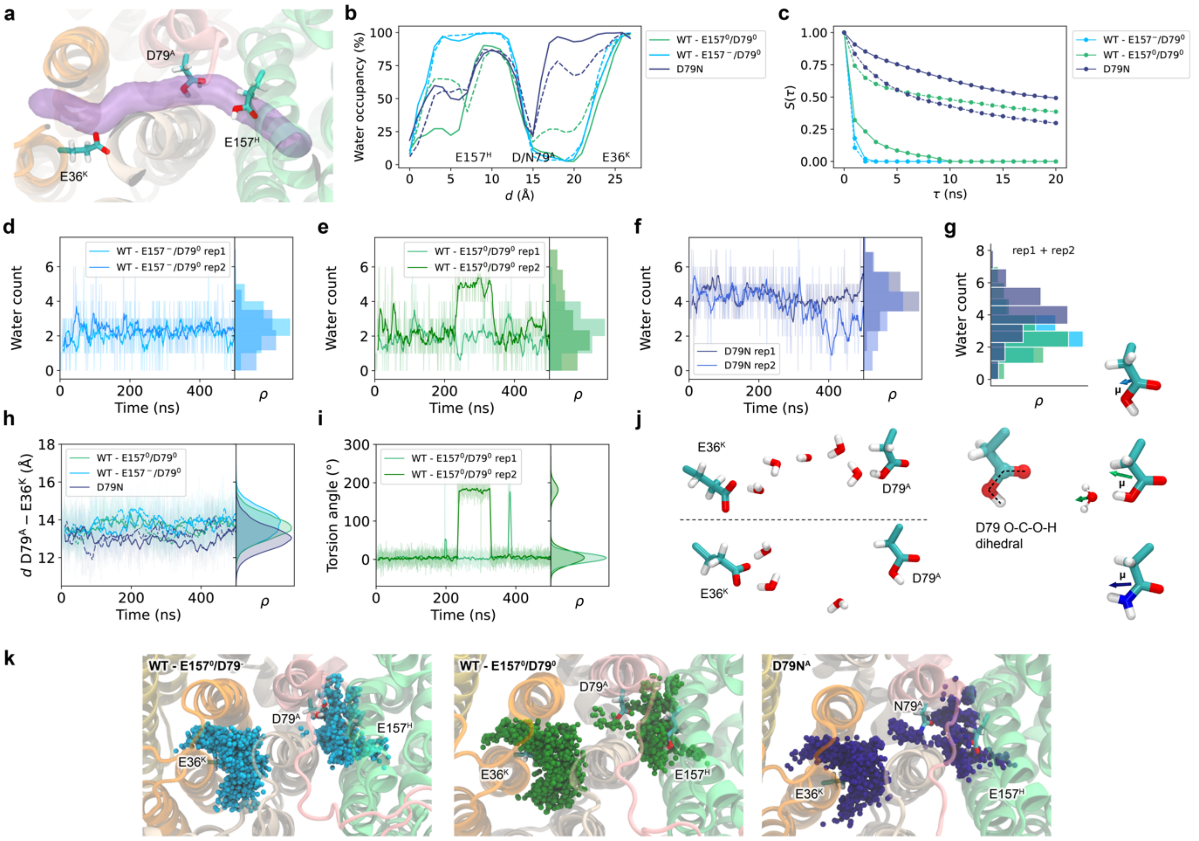


**Extended Data Fig. 3 | Hydration dynamics along the E-channel in Complex I and the D79N$^{A}$ variant. a,** Proton pathway along E157$^{H}$ via D79$^{A}$ to E36$^{K}$. **b,** Hydration level along the proton pathway. **c**, Survival probability of water molecules in the region between D79$^{A}$ and E36$^{K}$. **d-f**, Number of water molecules between E157$^{H}$/D79$^{A}$ and E36$^{K}$ during MD simulations (**d**, WT with E157$^{H}$ deprotonated; **e**, WT with E157$^{H}$ protonated; **f**: the D79N$^{A}$ variant). **g**, Histograms of the water count from **d-f**. **h**, Dynamics of the D(N)79$^{A}$–E36$^{K}$ distance during MD simulations. **i**, Dihedral angle of the protonated D79$^{A}$. The '*trans*' conformation correlates with higher hydration around the region (see panel **e**). **j**, Dihedral angle of D79$^{A}$ and dipole moment of D79$^{A}$/N79$^{A}$ in the different conformations. **k**, Top view of MD-averaged hydration of the proton pathway between E157$^{H}$/D79$^{A}$ and E36$^{K}$. *Left*: WT Complex I with Glu157$^{H}$ deprotonated, *middle*: WT Complex I with E157$^{H}$ protonated, *right:* the D79N$^{A}$ variant.

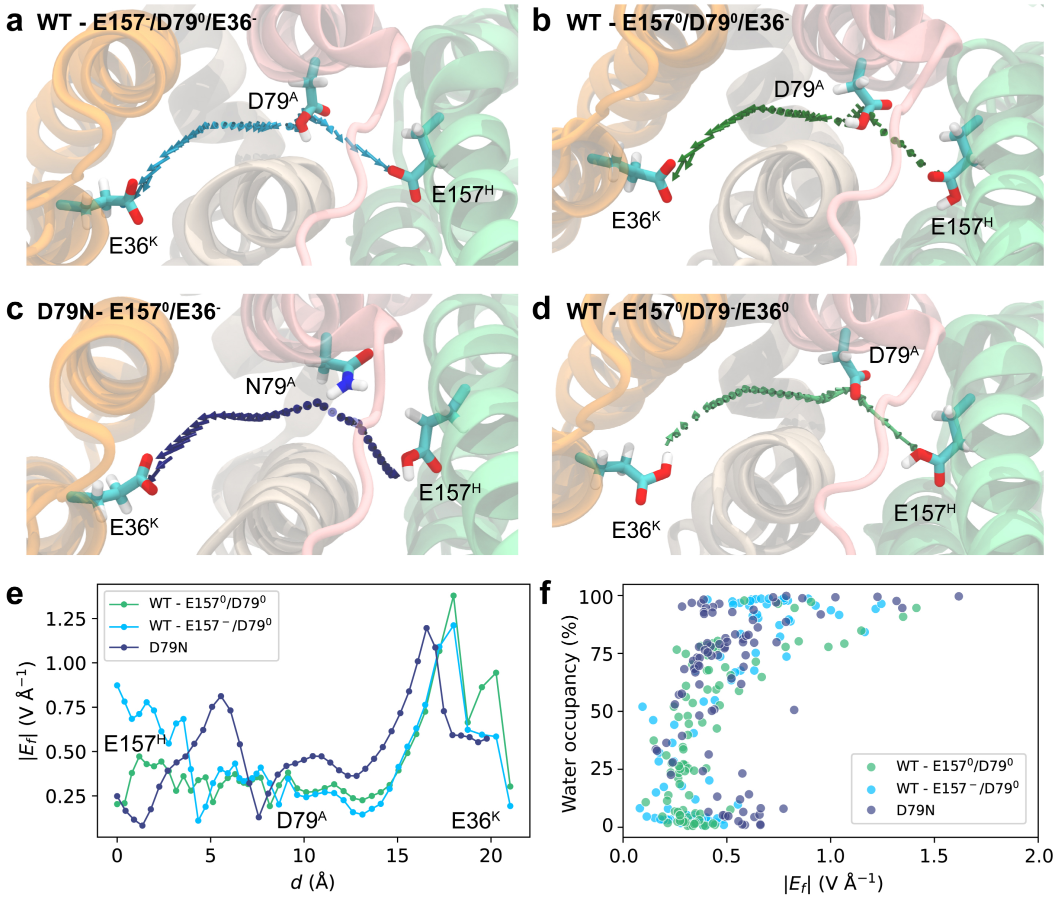


**Extended Data Fig. 4 | Electric field effects along the E-channel.** **a**, WT with E157$^{H}$ deprotonated; **b**, WT with E157$^{H}$ protonated; **c**, D79N$^{A}$ variant, and **d**, WT after proton transfer from D79$^{A}$ to E36$^{K}$, with E157$^{H}$ protonated. The average electric field ($E_f$) was computed from the classical MD simulations (average from 1 frame/ns). **e**, Electric field strength along the proton pathway. **f**, Correlation of the electric field strength and the water occupancy along the pathway.

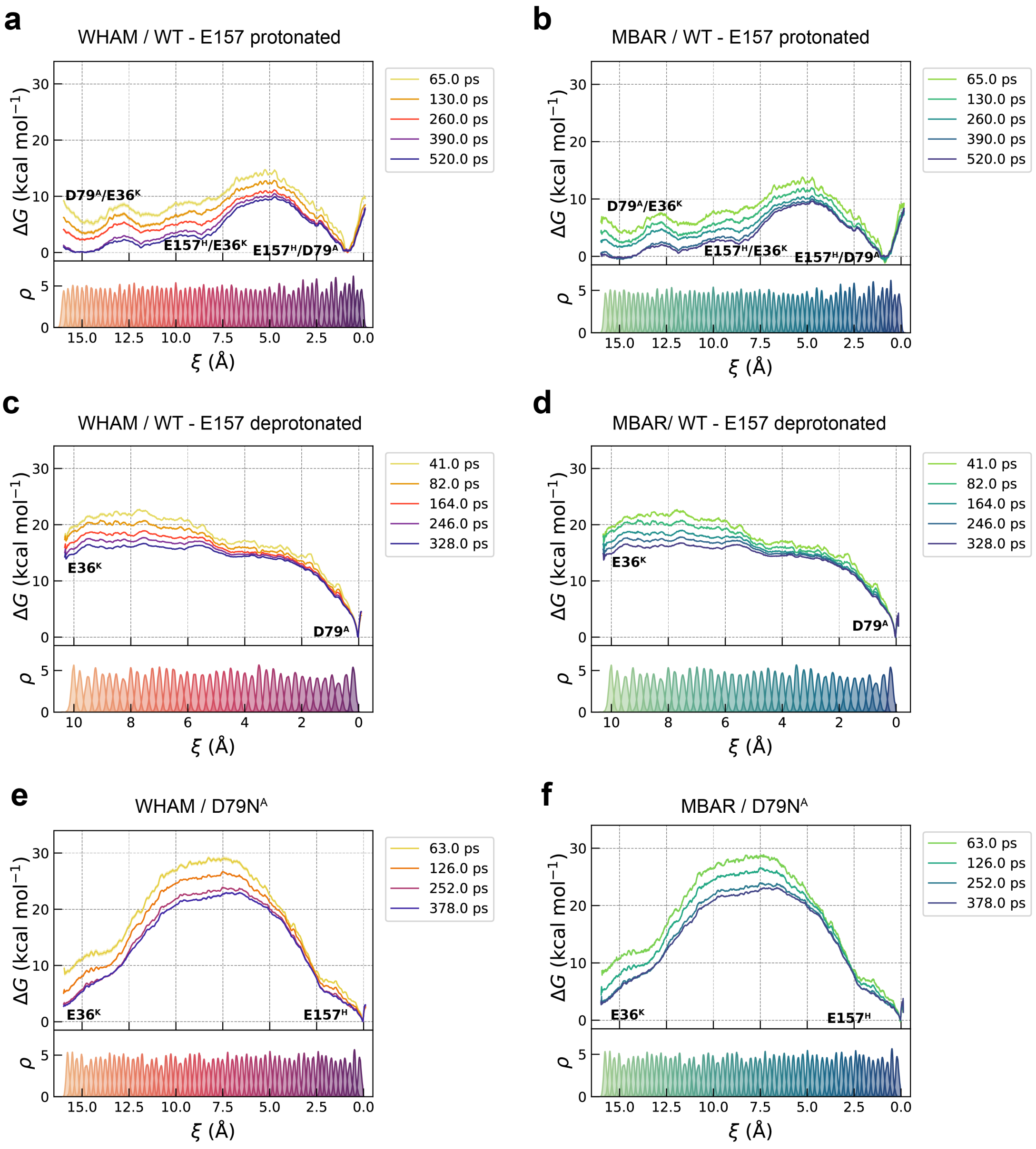


**Extended Data Fig. 5 | Convergence of the QM/MM free energy simulations.** Convergence of the free energy profiles at different levels of sampling (*top*) and histogram distributions of the QM/MM-US windows along the reaction coordinates (*bottom*) in **a**, WT with E157$^H$ protonated; **b**, WT with E157$^H$ deprotonated, and **c**, the D79N$^A$ variant.

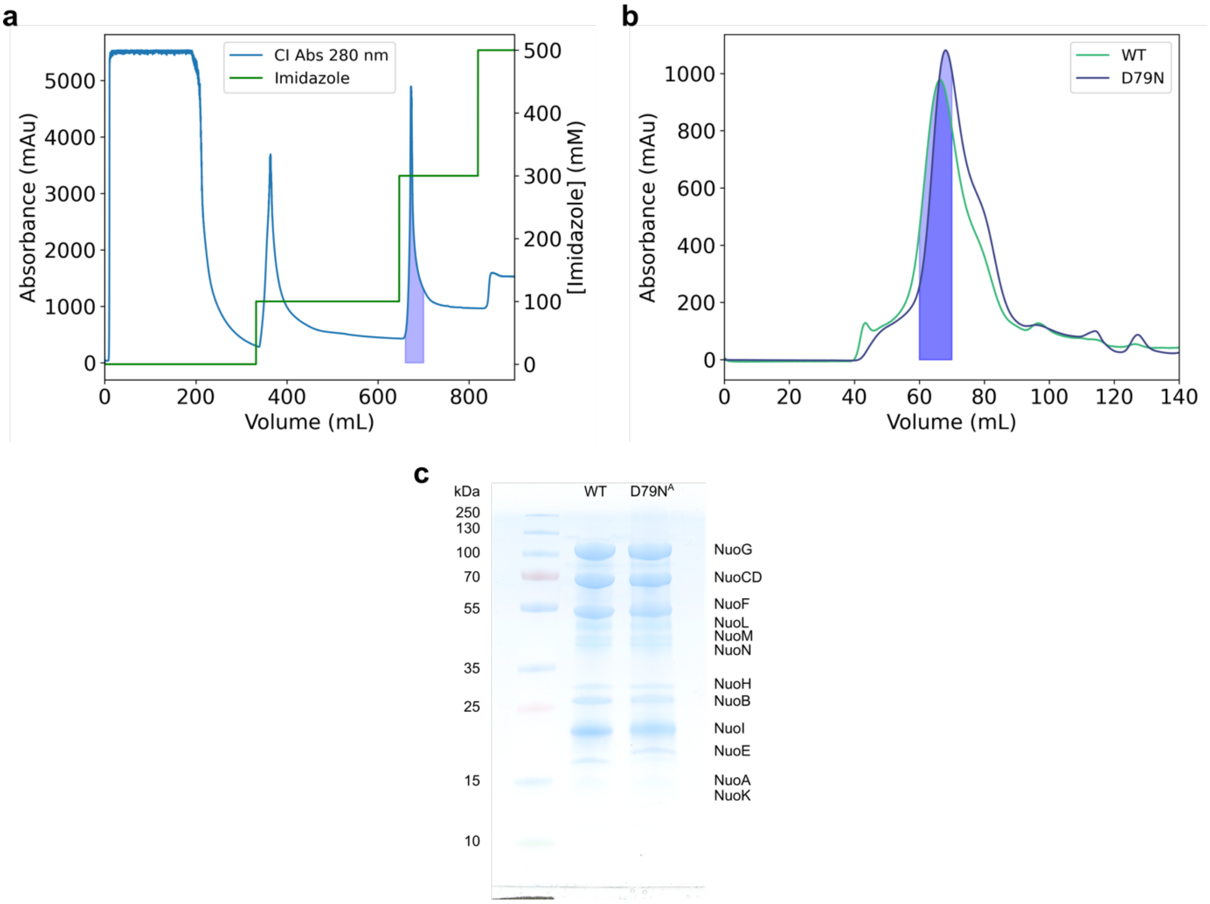


**Extended Data Fig. 6 | Protein purification**. **a**, Affinity chromatography profile monitored at 280 nm (*blue*) and imidazole concentration (*green*). Fractions collected upon elution with 300 mM imidazole are indicated by the integrated blue surface under the absorbance signal. **b**, Size exclusion chromatography profile monitored at 280 nm for Complex I (*green*) and the D79N$^{A}$ variant (*purple*). Collected fractions are shown by the integrated surface under the absorbance signal. **c**, SDS-PAGE fraction of purified Complex I and the D79N$^{A}$ variant.

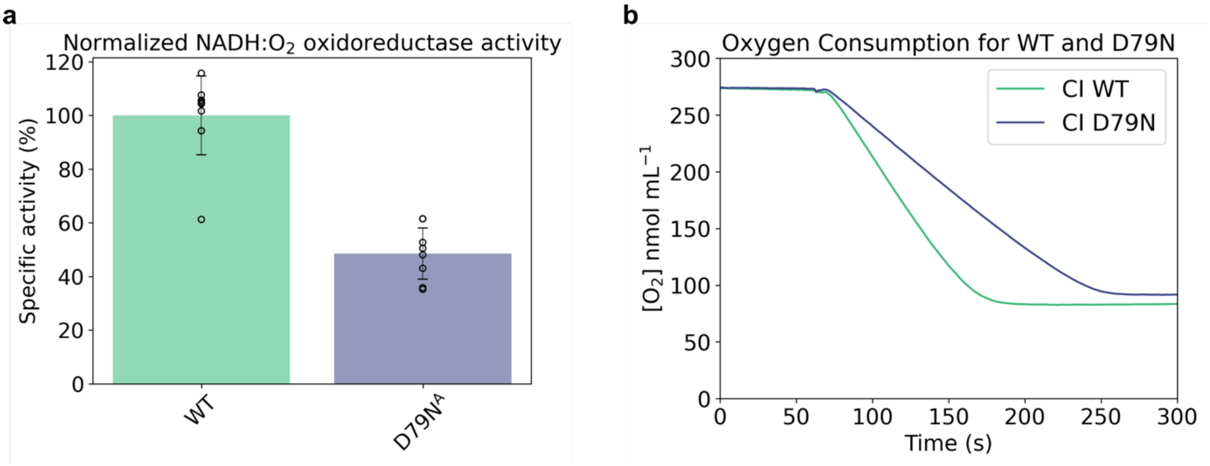


**Extended Data Fig. 7 | Oxygen consumption in cytoplasmic membranes. a**, Normalised oxygen consumption activity in cytoplasmic membranes from *E. coli* expressing the Complex I (100% activity corresponding to 0.55 ± 0.12 μmol $min^{-1}$ $mg^{-1}$) and the $D79N^A$ variant. **b**, Raw data of the oxygen consumption of membranes expressing Complex I and the $D79N^A$ variant (250-300 μg of protein).

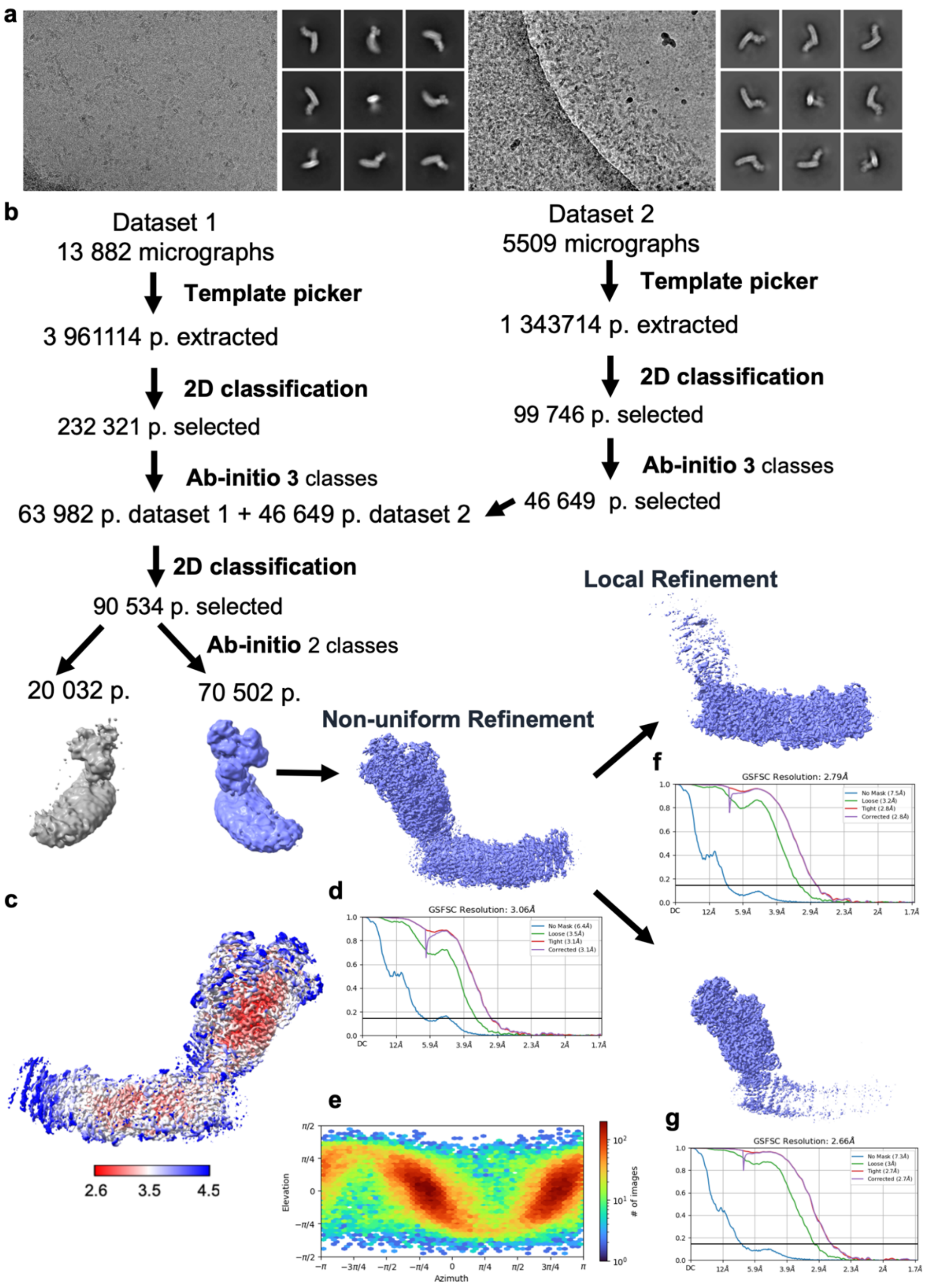


**Extended Data Fig. 8 | Cryo-EM data analysis and validation of Complex I**. **a**, Example of micrographs showing particle distribution for dataset 1 and 2 complemented with example of the 2D classification with various views. **b**, Schematic overview of data processing work flow. **c**, Local resolution representation of the non-uniform refined map of WT Complex I. **d**, Fourier shell correlation (FSC) curve of the non-uniform refined map of WT Complex I corrected for the effects of masking. **e**, Distribution of particle orientations of the non-uniform refinement of WT Complex I. **f**, Fourier shell correlation (FSC) curve of the local refinement of the membrane domain of WT Complex I corrected for the effects of masking. **g**, Fourier shell correlation (FSC) curve of the local refinement of the hydrophilic domain of WT Complex I, corrected for the effects of masking.

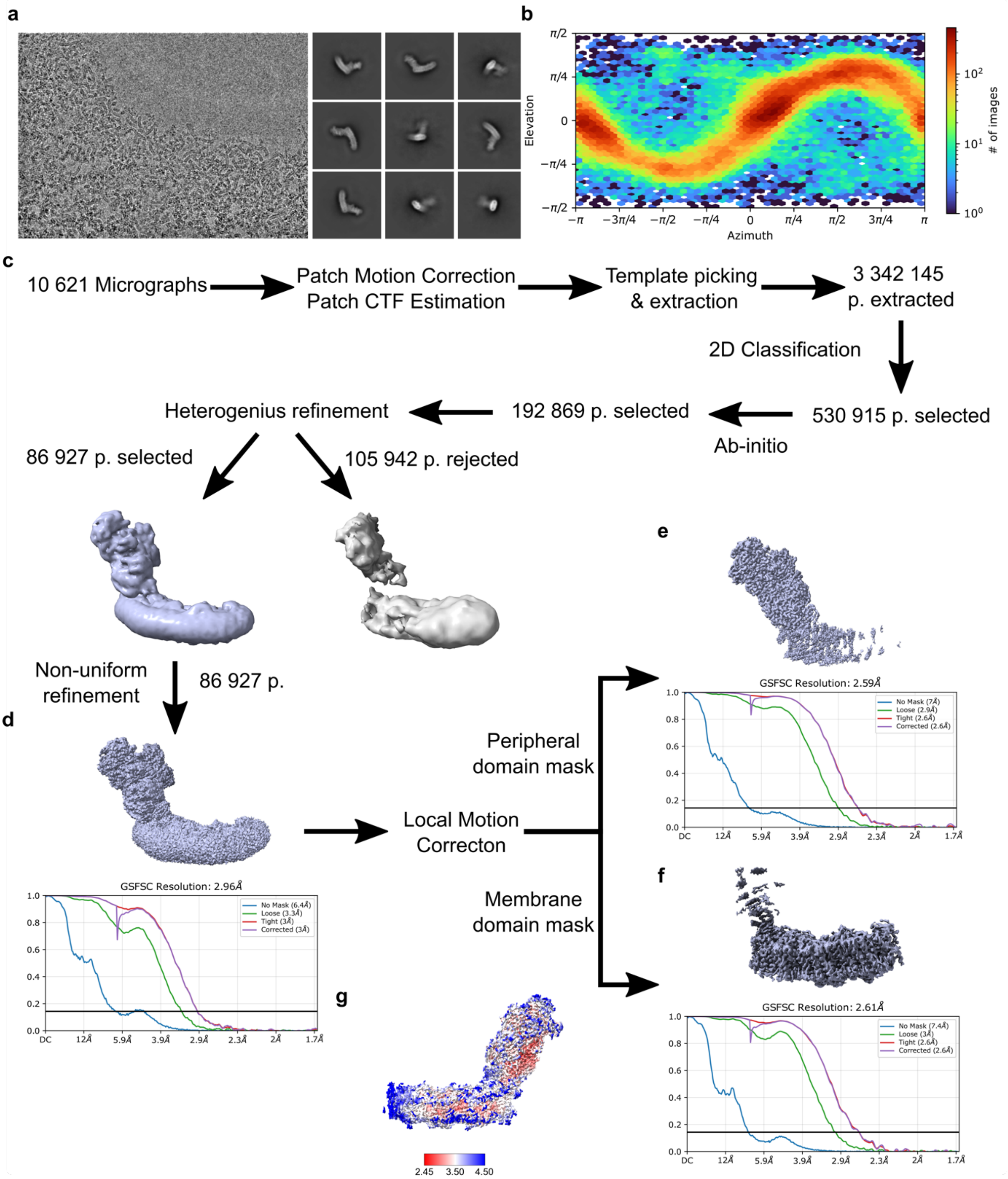


**Extended Data Fig. 9 | Cryo-EM data analysis and validation of the D79N^A variant. a**, Example of micrograph showing particle distribution and 2D classification in various views. **b**, Distribution of particle orientations of the non-uniform refinement of the D79N^A variant. **c**, Processing overview perform on CryoSparc v4.4.0. **d**, Fourier shell correlation (FSC) curve of the non-uniform refined map corrected for the effects of masking. **e**, Fourier shell correlation (FSC) curve of the local refinement of the membrane domain corrected for the effects of masking. **f**, Fourier shell correlation (FSC) curve of the local refinement of the hydrophilic domain, corrected for the effects of masking. **g**, Local resolution estimation representation of the non-uniform refined map.

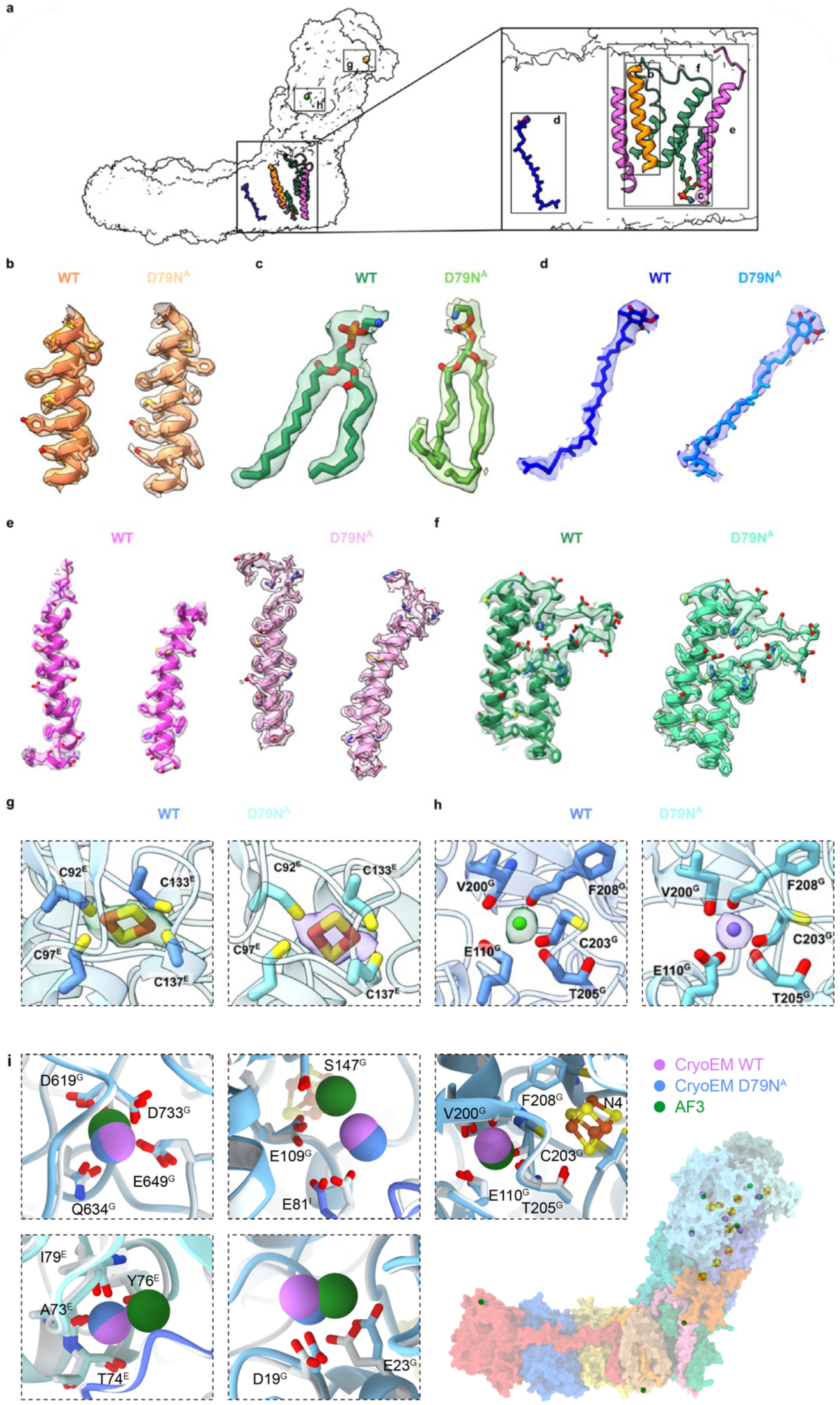


**Extended Data Fig. 10 | Example cryo-EM densities of key regions**. **a**, Location of regions shown in a-h. **b**, TM3^J. **c**, PE lipid (3PE) near NuoH. **d**, Membrane-bound ubiquinone-8 near NuoN. **e**, TM1-2 of NuoA, with an unresolved loop between residues Ser45^A and Asp55^A. **f**, TM5-6 loop of NuoH. **g**, FeS centre (N7) located in the NuoE subunit. **h**, $Ca^{2+}$ ion located in the NuoG subunit. **i**, Resolved $Ca^{2+}$ ions from cryo-EM structure (in *purple*), and based on AF3 prediction (in *green*).

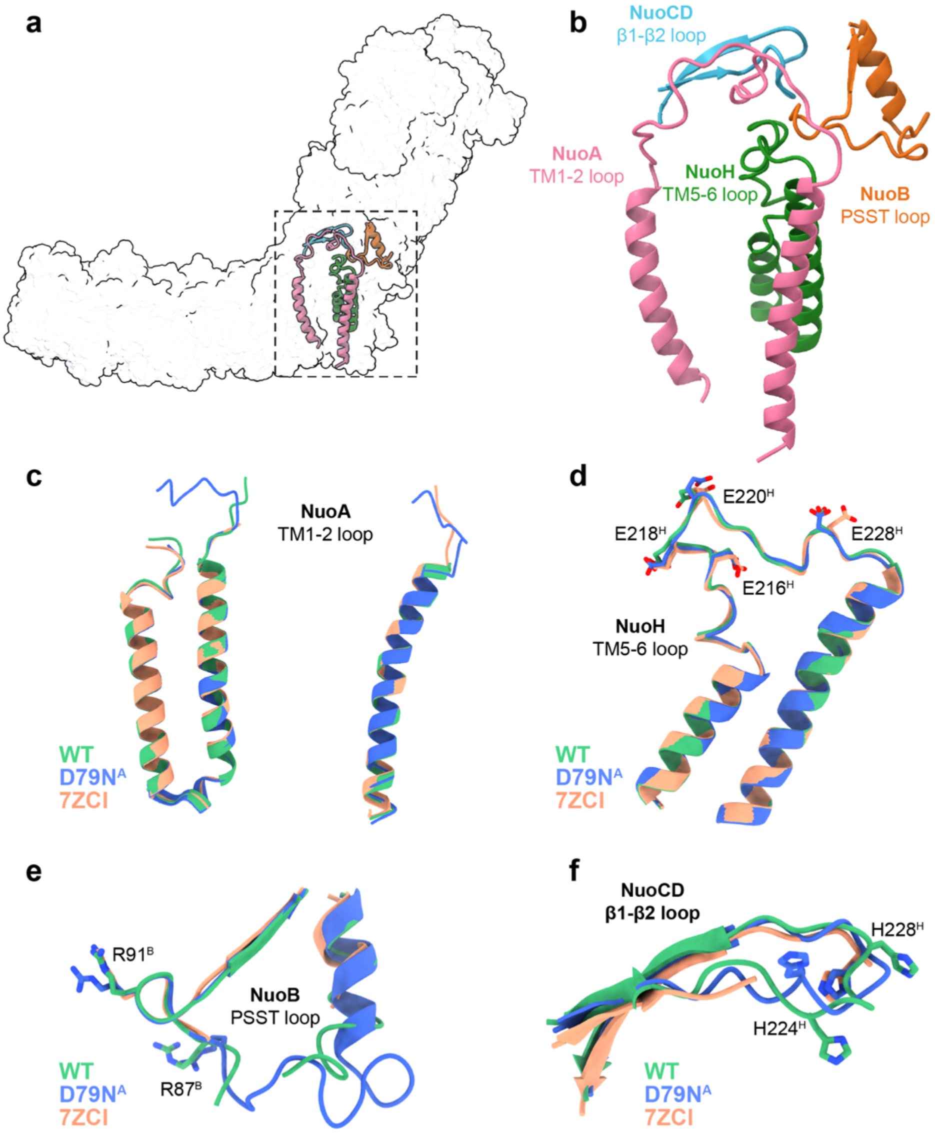


**Extended Data Fig. 11 | Structure of conserved loops. a**, Location of visualised regions. **b**, Closeup of the loops. Structure of **c**, TM1-2 of NuoA, **d**, TM5-6 loop of NuoH, **e**, PSST loop of NuoB, **f**, β1-β2 loop of NuoCD for Complex I and the D79N$^A$ variant, compared to a previous resting state structure of Complex I (PDB ID:7ZCI).

**Extended Data Table 1 | Activity of Complex I (WT) and the D79N[A] variant.**

| | Proton pumping activity | | | | |
|---|---|---|---|---|---|
| | ΔpH | | Δψ | | Orientation |
| | Rfu | % of WT | $\Delta Abs_{588-625}$ | % of WT | % |
| **WT** | 0.81 ± 0.06 | 100% ± 7.0% | 0.025 ± 0.003 | 100% ± 10.1% | 83.9% ± 7.4% |
| **D79N[A]** | 0.15 ± 0.02 | 18.5% ± 2.7% | 0.0056 ± 0.0009 | 22.3% ± 3.3% | 88.9% ± 2.6% |
| | **Oxidoreductase activity** | | | | |
| | **NADH:$O_2$ activity (in membranes)** | | | **NADH:DQ activity (in detergent)** | |
| | µmol $min^{-1}$ $mg^{-1}$ | % of WT | Normalised to FeCN | µmol $min^{-1}$ $mg^{-1}$ | % of WT |
| **WT** | 0.55 ± 0.12 | 100% ± 22.6% | 100% ± 14.7% | 28.8 ± 2.41 | 100% ± 8.35% |
| **D79N[A]** | 0.33 ± 0.08 | 59.9% ± 14.8 | 48.5% ± 9.5% | 7.57 ± 1.54 | 26.2% ± 5.35% |

**Extended Data Table 2 | List of designed primers.** Modified codons in bold, exchanged bases in italics. Restriction sites with silent mutations (in *italics*) are underlined.

| Oligonucleotide | Sequence |
|---|---|
| nuoA_D79N_fwd | 5'-CCATGTTCTTCGTTATCTTC***A*****AC**GTTGAAGC*TT*TGTATCTGTTCGCATGGTC-3' |
| nuoA_D79N_rev | 5'-GACCATGCGAACAGATACAAAGCTTCAACGTTGAAGATAACGAAGAACATGG-3' |
| seq_nuoA_D79N | 5'-GAGGTCGAAAAACGTG-3' |

### Extended Data Table 3 | List of employed buffers.

| | |
|---|---|
| **Cell resuspension buffer** | 50 mM MES pH 6.0, 50 mM KCl |
| **Membrane resuspension buffer** | 50 mM MES pH 6.0, 50 mM KCl, 5 mM $MgCl_2$, 10% glycerol |
| **IMAC buffer A** | 50 mM MES pH 6.0, 50 mM KCl, 5 mM $MgCl_2$, 10% glycerol, 0.005% LMNG, 20 mM Imidazole |
| **IMAC buffer B** | 50 mM MES pH 6.0, 50 mM KCl, 5 mM $MgCl_2$, 10% glycerol, 0.005% LMNG, 500 mM Imidazole |
| **SEC buffer C** | 50 mM MES pH 6.0, 50 mM KCl, 5 mM $MgCl_2$, 10% glycerol, 0.005% LMNG |
| **Cryo-EM buffer** | 50 mM MES pH 6.0, 150 mM KCl, 5 mM $MgCl_2$, 2% glycerol, 0.005% LMNG |
| **Reconstitution buffer** | 50 mM MES pH 6.7, 150 mM NaCl |
| **FeCN buffer** | 50 mM MES pH 6.7, 150 mM NaCl, 1 mM $K_3Fe(CN)_6$ |
| **ACMA buffer** | 50 mM MES pH 6.7, 150 mM NaCl, 4 µM ACMA |
| **Oxonol VI buffer** | 50 mM MES pH 6.7, 150 mM NaCl, 5 µM oxonol, 100 nM monensin, 300 mM mannitol |

### Extended Data Table 4 | CryoEM data collection, refinement, and validation statistics.

| **Data collection, processing** | **Complex I overall** PDB ID: 9TAJ | **Complex I membrane** PDB ID: 9TAK | **Complex I hydrophilic** PDB ID: 9TAL | **D79N[A] variant overall** PDB ID: 9TAM | **D79N[A] variant membrane** PDB ID: 9TAO | **D79N[A] variant hydrophilic** PDB ID: 9TAN |
|---|---|---|---|---|---|---|
| Voltage (kV) | 300 | 300 | 300 | 300 | 300 | 300 |
| Magnification | 105,000 | 105,000 | 105,000 | 105,000 | 105,000 | 105,000 |
| Electron exposure ($e^-/Å^2$) | 40 | 40 | 40 | 40 | 40 | 40 |
| Pixel size (Å) | 0.825 | 0.825 | 0.825 | 0.825 | 0.825 | 0.825 |
| Defocus range (µm) | -2.0 to -0.6 | -2.0 to -0.6 | -2.0 to -0.6 | -2.0 to -0.6 | -2.0 to -0.6 | -2.0 to -0.6 |
| Defocus step (µm) | -0.2 | -0.2 | -0.2 | -0.2 | -0.2 | -0.2 |
| Symmetry imposed | None (C1) | None (C1) | None (C1) | None (C1) | None (C1) | None (C1) |
| Initial particle images (number) | 3,961,114 + 1,343,714 | | | 3,342,145 | | |
| Final particle images (number) | 70502 | | | 146537 | | |
| FSC threshold | 0.143 | 0.143 | 0.143 | 0.143 | 0.143 | 0.143 |
| Map resolution (Å) | 3.06 | 2.79 | 2.66 | 2.93 | 2.61 | 2.59 |
| **Refinement** | | | | | | |
| CC (mask) | 0.8 | 0.92 | 0.93 | 0.63 | 0.71 | 0.85 |
| Resolution estimates (Å) | | | | | | |
| d 99 | | | | | | |
| Masked | 3.7 | 3.4 | 3.2 | 3.2 | 3.3 | 3.2 |
| Unmasked | 3.5 | 3.2 | 3.1 | 3.1 | 3.1 | 3.0 |
| d FSC model, 0/0.143/0.5 (Å) | | | | | | |
| Masked | 3.0/3.0/3.4 | 2.7/2.8/3.0 | 2.6/2.6/2.8 | 2.9/2.9/3.6 | 2.5/2.6/3.1 | 2.5/2.6/2.8 |
| Unmasked | 3.0/3.0/3.8 | 2.8/2.8/3.3 | 2.6/2.7/3.2 | 2.9/3.1/4.1 | 2.5/2.7/3.4 | 2.6/2.6/3.0 |
| Model composition | | | | | | |
| Protein residues | 4698 | 2358 | 2552 | 4717 | 2276 | 2449 |
| Non-hydrogen atoms | 38186 | 19437 | 20425 | 38164 | 18397 | 19678 |
| Ligands | 39 | 24 | 15 | 32 | 20 | 14 |
| Water | 428 | 166 | 258 | 416 | 184 | 296 |
| **B factors ($Å^2$)** | | | | | | |
| | Min/max/mean | | | | | |
| Protein residues | 19.42/170.42/59.36 | 29.16/207.45/66.59 | 19.42/185.48/59.79 | 0.01/130.10/22.09 | 0.01/130.10/21.75 | 0.64/110.12/22.92 |
| Ligands | 30.00/175.57/59.36 | 37.72/175.57/88.66 | 30.00/73.33/48.65 | 1.32/93.30/43.00 | 8.29/79.35/44.14 | 1.32/93.30/35.91 |
| water | 29.39/104.02/55.75 | 40.80/104.02/67.30 | 29.39/93.14/48.44 | 0.07/78.52/17.44 | 0.07/65/55/20/09 | 0.91/78.52/14.43 |
| **Validation** | | | | | | |
| Ramachandran plot | | | | | | |
| Favoured (%) | 96.12 | 95.67 | 96.24 | 95.94 | 96.12 | 96.26 |
| Allowed (%) | 3.84 | 4.29 | 3.76 | 4.04 | 3.84 | 3.70 |
| Disallowed (%) | 0.04 | 0.04 | 0.00 | 0.02 | 0.04 | 0.04 |

**Extended Data Table 5 | List of MD simulations.**

| Simulation | System | E-channel state | Started from | Time (ns) |
|---|---|---|---|---|
| **S1/S2** | WT | E157$^-$/**D79$^0$**/E36$^-$/E72$^-$ | - | 2×500 |
| **S3/S4** | WT | **E157$^0$**/**D79$^0$**/E36$^-$/E72$^-$ | - | 2×500 |
| **S5/S6** | D79N[A] | **E157$^0$**/ E36$^-$/E72$^-$ | - | 2×500 |
| **S7/S8** | WT | E157$^-$/D79$^-$/**E36$^0$**/E72$^-$ | S1/S2 | 2×100 |
| **S9/S10** | WT | **E157$^0$**/D79$^-$/**E36$^0$**/E72$^-$ | S3/S4 | 2×100 |
| **S11/S12** | WT | E157$^-$/**D79$^0$**/**E36$^0$**/E72$^-$ | S9/S10 | 2×100 |
| **S13/S14** | WT | E157$^-$/D79$^-$/E36$^-$/**E72$^0$** | S7/S8 | 2x100 |
| **S15/S16** | WT | **E157$^0$**/D79$^-$/E36$^-$/E72$^-$ | - | 2x100 |
| **S17/S18** | D79N[A] | E157$^-$/ E36$^-$/E72$^-$ | - | 2x100 |
| | | | **Total** | 3.6 μs |

**Extended Data Table 6 | List of QM/MM simulations.**

| Simulation | System | E-channel state | QM region | Total sampling (ps) |
|---|---|---|---|---|
| **Q1** | WT | $E157^0/D79^{0-}$ | $D79^A$, $E36^K$, $E157^H$, $F75^A$, $V76^A$, $T153^H$, $Y156^H$, $V58^J$, $Y59^J$, $A62^J$, $N40^K$, $A73^K$, 24 waters | 65 × (8 ps) = 520 ps |
| **Q2** | WT | $E157^-/D79^0$ | $D79^A$, $E36^K$, $E157^H$, $F75^A$, $V76^A$, $T153^H$, $Y156^H$, $V58^J$, $Y59^J$, $A62^J$, $N40^K$, $A73^K$, 23 waters | 41 × (8 ps) = 328 ps |
| **Q3** | $D79N^A$ | $E157^{0-}$ | $N79^A$, $E36^K$, $E157^H$, $F75^A$, $V76^A$, $T153^H$, $Y156^H$, $V58^J$, $Y59^J$, $A62^J$, $N40^K$, $A73^K$, 26 waters | 63 × (6 ps) = 378 ps |
| | | | **Total:** | **1226 ps** |

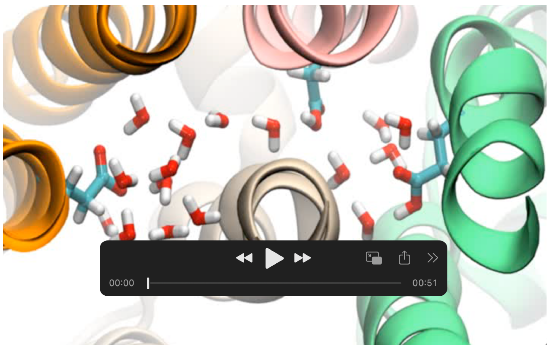


**Extended Data Movie 1 |** Proton transfer along the E-channel of Complex I.

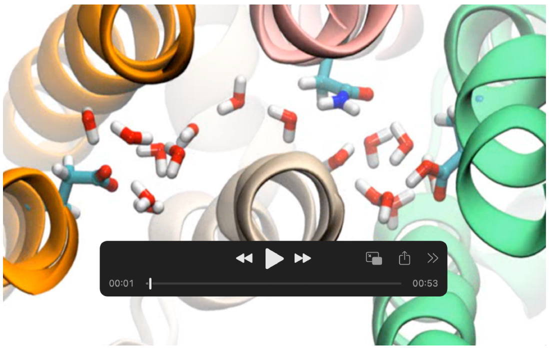


**Extended Data Movie 2 |** Proton transfer along the E-channel of the D79N$^{A}$ variant.
